# Ubiquitination is essential for recovery of cellular activities following heat shock

**DOI:** 10.1101/2021.04.22.440934

**Authors:** Brian A. Maxwell, Youngdae Gwon, Ashutosh Mishra, Junmin Peng, Ke Zhang, Hong Joo Kim, J. Paul Taylor

**Affiliations:** Department of Cell and Molecular Biology, St. Jude Children’s Research Hospital, Memphis, TN, USA; Department of Structural Biology Department, St. Jude Children’s Research Hospital, Memphis, TN, USA; Department of Neuroscience, Mayo Clinic Florida, Jacksonville, FL 32224; Howard Hughes Medical Institute, Chevy Chase, MD 20815

## Abstract

Eukaryotic cells respond to stress via adaptive programs that include reversible shutdown of key cellular processes, the formation of stress granules, and a global increase in ubiquitination. The primary function of this ubiquitination is generally considered to be tagging damaged or misfolded proteins for degradation. Here we show that different types of stress generate distinct ubiquitination patterns. For heat stress, ubiquitination correlates with cellular activities that are downregulated during stress, including nucleocytoplasmic transport and translation, as well as with stress granule constituents. Ubiquitination is not required for the shutdown of these processes or for stress granule formation, but is essential for resumption of cellular activities and for stress granule disassembly. These findings indicate that stress-induced ubiquitination primes the cell for recovery following heat stress.

**One Sentence Summary:** Stress-induced ubiquitination is essential for recovery of cellular activities following heat stress.

## Main Text

Eukaryotic cells have a stereotypical response to a variety of stresses that aids in their survival until normal growth conditions are restored (*1, 2*). This stress response is associated with inhibition of global translation along with upregulated expression of select stress response factors (*1–7*). In addition to programmed shutdown of translation, evidence suggests that other biological pathways are also shut down or perturbed, including nucleocytoplasmic transport (*8–13*), RNA splicing (*14, 15*), and cell cycle activities (*16–18*), among others. Inhibition of translation and subsequent polysome disassembly leads a rise in the cytoplasmic concentration of free mRNA that culminates in the formation of cytoplasmic condensates known as stress granules (*19*). These stress-induced cellular changes are adaptive in the short term but require coordinated reversal after stress is removed in order to resume cellular activities and reestablish homeostasis. Accordingly, upon the removal of stress, stress granules disassemble while translation and other biological pathways return to normal activity levels on a time scale of minutes to a few hours (*2, 20–22*). However, the mechanisms that facilitate this recovery are not well understood.

One hallmark of the cellular response to many types of stress is a global increase in poly-ubiquitin conjugation, which has largely been attributed to increased protein quality control (PQC) activity in response to stress-induced protein damage and translation arrest (*1–3*). This increased activity is required for the ubiquitin-dependent degradation of both defective ribosomal products (DRiPs), the primary source of misfolded proteins in cells under normal conditions (*4*), as well as proteins susceptible to stress-induced misfolding, such as thermolabile proteins or proteins with intrinsically disordered domains (*2, 5–7*). Interestingly, immunostaining and proteomics have demonstrated the presence of ubiquitin and some ubiquitin pathway proteins in stress granules (*23–28*), and studies have shown that several deubiquitinating enzymes can contribute to regulating stress granule dynamics (*23, 27*), although the mechanism of this regulation is largely unknown. Furthermore, when PQC function is disrupted, ubiquitinated DRiPs can aberrantly accumulate in stress granules and impair stress granule function (*29–31*). Several investigations have examined the possibility that ubiquitination may play an additional role in regulating stress granule dynamics, yielding conflicting results: whereas one study reported that the ubiquitin-selective segregase VCP facilitates disassembly of arsenite-induced stress granules (*29*), a more recent study found minimal ubiquitination of stress granule proteins and no detectable role for ubiquitin conjugation in the formation or disassembly of arsenite-induced stress granules (*32*). The role of ubiquitination in regulating stress granule dynamics remains unresolved, as does the role of ubiquitination in regulating other cellular activities that are altered or shut down by stress.

Analysis of stress-dependent changes to the global protein ubiquitin modification landscape, or “ubiquitinome,” can provide unbiased insights into details of the stress response by connecting differentially ubiquitinated proteins with specific biological pathways. To this end, several proteomic techniques have been developed to allow for quantitative comparisons of ubiquitinomes from differentially treated cell or tissue samples (*33–36*). Analysis of the ubiquitinome in yeast has suggested that heat shock-induced ubiquitination primarily targets cytosolic misfolded proteins (*37, 38*), consistent with the presumption that ubiquitination in this setting relates exclusively to PQC, though it remains unknown whether the same holds true in human cells. Furthermore, comprehensive analysis to directly compare changes in ubiquitination in response to various stresses has not been undertaken, and it remains unclear to what extent ubiquitination is similar or different in these different contexts.

Here we used a series of complementary proteomic analyses in mammalian cells, including tandem ubiquitin binding entity (TUBE) enrichment (*35*), di-GLY enrichment (*36*), and tandem mass tagging (TMT) (*39*), to quantitatively assess the stress-responsive ubiquitination elicited by five different types of stress. We determined that each type of stress is associated with unique but overlapping patterns of ubiquitination. Taking a deep dive into the heat stress response, we determined that in addition to playing a role in PQC, ubiquitination also correlates with biological processes that are downregulated or shut off. Remarkably, ubiquitination is not involved in shutting off these processes, but is essential for the restoration of normal activity during the recovery phase following stress, including resumption of nucleocytoplasmic transport and translation, and the disassembly of stress granules.

## Results

### Different cellular stresses induce distinct patterns of ubiquitination

To investigate whether cells deploy different patterns of ubiquitination to cope with different types of stress, we performed proteomic analysis following TUBE-based capture of the ubiquitinome in HEK293T cells exposed to heat stress (43°C), oxidative stress (sodium arsenite), osmotic stress (sorbitol), UV stress, or proteasomal inhibition (bortezomib) (**Fig. 1, A and B**). Following exposure to each type of stress, lysates were incubated with HALO-linked TUBE resin to capture ubiquitin conjugates, followed by label-free LC-MS/MS (**Fig. 1, A and B**). Immunoblotting for poly-ubiquitin indicated that all stress types except UV led to an increase in total poly-ubiquitin conjugates (**input blot, Fig. 1C**) and that the TUBEs captured the vast majority of the ubiquitinated material (**compare bound and unbound blots, Fig. 1C**). We used harsh buffer conditions in our TUBE protocol to minimize non-covalent binding of non-ubiquitinated proteins to poly-ubiquitin chains or ubiquitin conjugates. We detected ~200-300 ubiquitinated proteins per stress condition and a total of 500 unique proteins across all five stress conditions, many of which increased or decreased in abundance in response to stress (**Fig. 1D, table S1**).

**Fig. 1.**
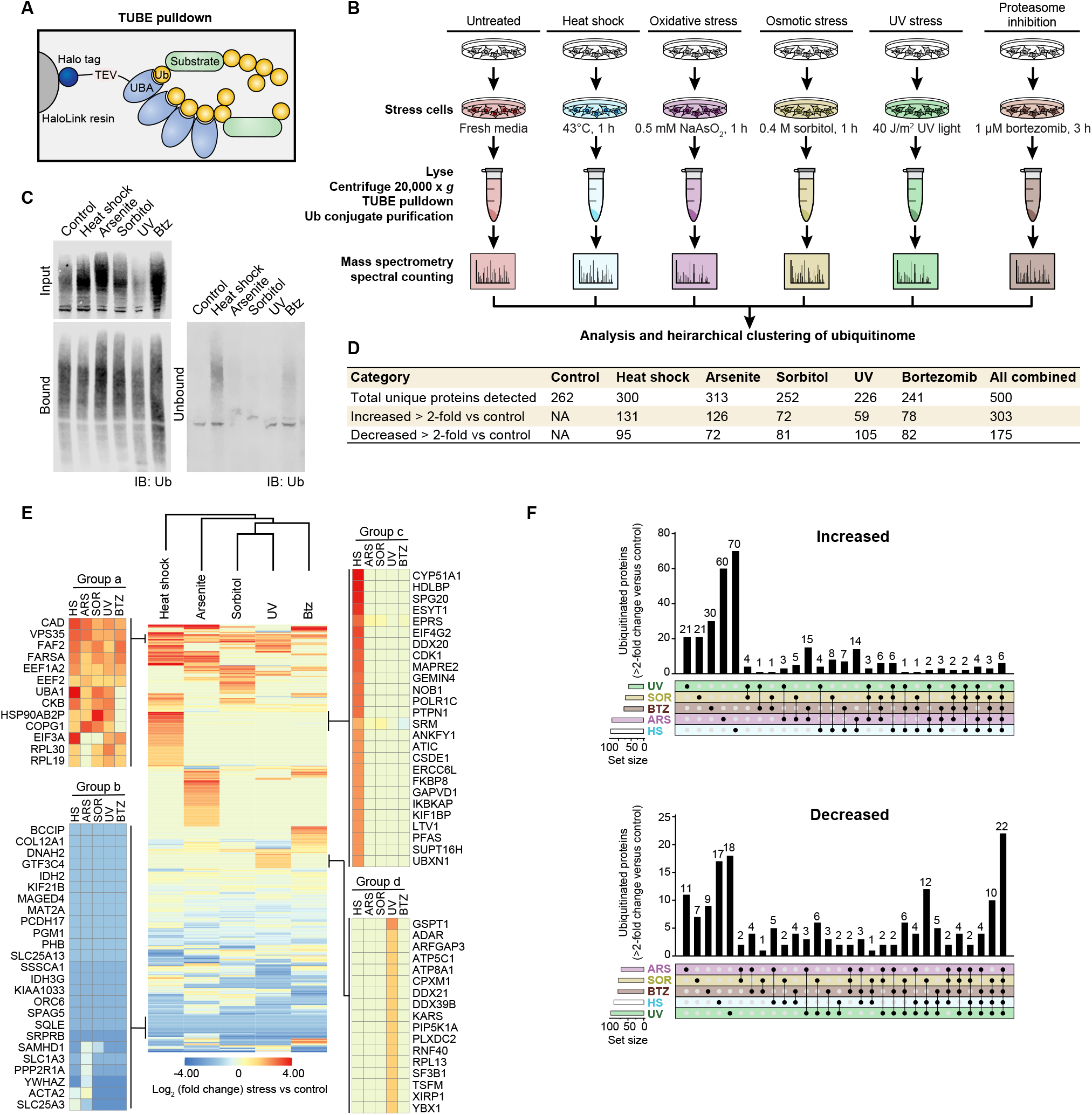
Different cellular stresses induce different patterns of ubiquitination. (**A**) Schematic representation of HALO-linked TUBE resin and ubiquitinated protein capture. (**B**) Workflow for sample preparation and TUBE proteomics. HEK293T cells were treated with the indicated stress or incubated with fresh media as a control, then lysed and ubiquitinated proteins were captured with TUBEs prior to MS spectral counting. (**C**) Immunoblot showing TUBE capture of ubiquitinated proteins from cells stressed as in (B). Indicated fractions were immunoblotted with poly-ubiquitin antibody PD41. (**D**) Summary of statistics from proteomic analysis. (**E**) Heatmap illustrating global stress-induced changes to the ubiquitinome. Colors indicate log_2_ fold change in spectral counts of stressed samples versus control samples. Example groups of proteins with a shared pattern are shown in the blow-ups. (**F**) UpSet plot indicating proteins changed > 2-fold versus control for each stress.

In comparing the ubiquitin-conjugate landscape across the five modes of stress, unsupervised hierarchical clustering revealed that each type of stress was associated with a distinct pattern of altered ubiquitination (**Fig. 1E**). Indeed, the majority of proteins with increased or decreased ubiquitination were associated with a single type of stress (**Fig. 1F).** Several patterns emerged from this analysis. Some proteins showed increased or decreased ubiquitination in response to all 5 stresses, suggesting that certain ubiquitination events are part of a non-specific, generalized stress response (**e.g., Groups a and b, Fig. 1E**). On the other hand, some prominent groups of proteins showed altered ubiquitination in a stress-specific manner. For example, ubiquitination of some proteins was increased exclusively in response to heat shock (**Group c, Fig. 1E**) or UV exposure (**Group d, Fig. 1E**). The emergence of these distinct protein groups suggested the existence of stress-specific patterns of ubiquitination that represent distinct adaptive responses to distinct stressors. To test this hypothesis, we deeply investigated stress-responsive ubiquitination in response to heat stress and oxidative stress, stresses that evoked a large number of unique changes in ubiquitination. Whereas the cellular response to these stresses have been intensely studied, insight into stress-responsive ubiquitination has been very limited and largely assumed to reflect non-selective degradation of misfolded or damaged proteins.

### Heat stress leads to rapid increases in total poly-ubiquitination and a time-dependent shift in solubility of ubiquitin conjugates

We first examined the kinetics of poly-ubiquitin conjugate accumulation during heat shock and recovery. Heat stress-induced ubiquitination occurred rapidly, with increased poly-ubiquitin conjugates apparent as early as 15 minutes and peaking within 30 minutes following heat shock (**Fig. 2A**). The accumulated poly-ubiquitin conjugates remained at a constant elevated level during prolonged stress for at least 150 minutes (**Fig. 2A**). Prolonged heat shock has been reported to result in progressive loss of solubility in a fraction of the proteome (*40–43*), although the relationship to ubiquitination has not been examined previously. Here we observed a time-dependent dynamic shift in the ubiquitinome between soluble and insoluble fractions. Specifically, accumulation of poly-ubiquitin conjugates in the soluble fraction peaked at about 30 minutes and then progressively declined over the course of the experiment. In contrast, poly-ubiquitin conjugates accrued more slowly in the insoluble fraction, reaching a plateau after 60 minutes (**Fig. 2A**). The accumulation of poly-ubiquitin conjugates after heat stress was temporary, returning to baseline within 3 hours after a 30-minute heat shock (**Fig. 2B**).

**Fig. 2.**
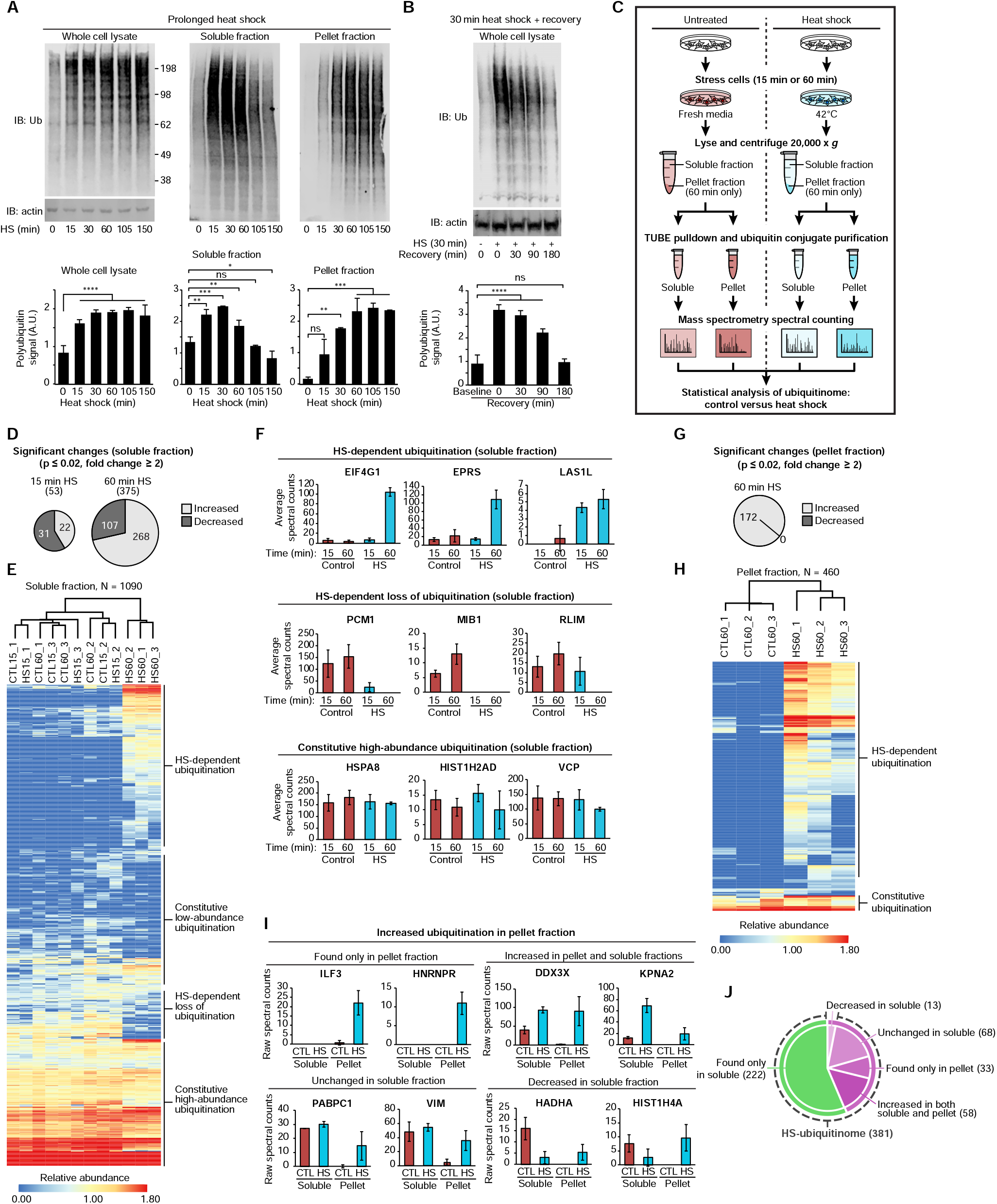
Heat shock induces significant changes to the global ubiquitinome. (**A-B**) Immunoblots showing poly-ubiquitination levels at various time points of heat shock (A) and during recovery from 30 min heat shock (B) in whole cell lysates, soluble fractions, and pellet fractions after centrifugation at 14,000 × *g*. Bar graphs indicate quantification of Western blots from ≥ 3 replicates for each experiment. Error bars indicate s.d. ns, not significant; \**P* < 0.05, \*\**P* < 0.01, \*\*\**P* < 0.001, \*\*\*\**P* < 0.0001 ANOVA with Dunnett’s test. (**C**) Workflow for soluble and pellet fraction sample preparation and TUBE proteomic analysis. (**D**) Charts indicating the number of significantly changed proteins in soluble fractions for heat-shocked samples (15 min or 60 min) compared to control. (**E**) Heatmap illustrating heat shock-induced changes to the global ubiquitinome in soluble fractions. Colors indicate relative abundance as quantified by spectral counting. (**F**) Spectral count values for several representative proteins of indicated categories; averages are shown from three replicates of each sample condition in soluble fractions. Error bars indicate s.d. (**G**) Chart indicating the number of significantly changed proteins in pellet fractions for heat-shock samples compared to control. (**H**) Heatmap illustrating heat shock-induced changes to the global ubiquitinome in pellet fractions. (**I**) Spectral count values for several representative proteins of indicated categories; averages are shown from three replicates of each sample condition in soluble and pellet fractions. CTL, control; HS, 60 min heat shock. Error bars indicate s.d. (**J**) Chart indicating the number of proteins with heat shock-dependent increase in ubiquitination as detected in soluble and/or pellet fractions. Proteins detected in the pellet fraction are shown in purple, proteins detected only in the soluble fraction are shown in green. Dashed line indicates categories included in the heat shock ubiquitinome.

With this kinetic profile in mind, we next performed deep proteomic analysis of the heat stress-associated ubiquitinome by TUBE-LC-MS/MS. We examined this ubiquitinome in the soluble fraction at an early time point (15 min) corresponding to the initial increase in poly-ubiquitination, and at a later time point (60 min) in both soluble and pellet fractions (**Fig. 2C**). This quantification of over 1,000 ubiquitinated proteins in the TUBE-recovered samples indicated good depth of coverage and reproducibility (**fig. S1, A and B, table S2**). We also verified heat stress-induced increase in poly-ubiquitination of seven representative proteins by immunoblotting (**fig. S1C**). Importantly, replicating this experiment in U2OS cells gave results very similar to those in HEK293T cells, indicating that heat stress-induced changes in ubiquitination are not specific to one cell type (**fig. S1D and table S3**).

In examining the temporal profile of changes in the ubiquitinome in the soluble fraction, we observed only modest changes at 15 minutes but more substantial changes at 60 minutes following heat stress (**Fig. 2D,-F**). These results indicate that despite an increase in a poly-ubiquitin smear detected by immunoblotting at 15 minutes (**Fig. 2A**), the global shift in the ubiquitinated protein landscape is better reflected later in the stress response process. We therefore directed the majority of our subsequent analyses to examining changes to the ubiquitinome after 60 minutes of heat shock.

Significant changes to the ubiquitinome were also detected in the pellet fraction. Whereas we detected no loss of ubiquitinated proteins in the pellet fraction, there was a significant increase in 172 ubiquitinated proteins after 60 minutes of heat stress (**Fig. 2, G and H**). Roughly 80% of these proteins had also been detected in the soluble fraction, where these proteins had been found to be enriched, depleted, or unchanged in response to heat stress (**Fig. 2, I and J**). Importantly, our analysis indicated that while some ubiquitinated proteins do accumulate in the pellet fraction following heat shock, an individual protein’s ubiquitination status was not predictive of its solubility. Therefore, in subsequent analyses we assessed the significance of heat stress-induced ubiquitination of all proteins irrespective of whether they appeared in the soluble fraction, pellet fraction, or both. Altogether, our analysis demonstrated that heat stress induced a global shift in the ubiquitin landscape, resulting in a “heat shock ubiquitinome” of 381 proteins which we define as all proteins showing a predicted net total increase in ubiquitination in response to heat stress (**Fig. 2J and table S4**).

### Heat and arsenite stress induce different changes to the ubiquitinome

In the initial characterization of ubiquitination patterns that arise in response to different stressors, unsupervised hierarchical clustering showed that the pattern of ubiquitination following arsenite stress was the most similar to that generated by heat stress (**Fig. 1E**). To examine this more deeply, we characterized arsenite stress-induced ubiquitination at 60 minutes in triplicate following the same approach that we used to define heat stress-induced ubiquitination. After identifying proteins whose ubiquitination was significantly altered relative to control cells (**tables S4 and S5**), we compared the responses to arsenite stress versus heat stress. Consistent with our pilot study, this deep analysis confirmed significant overlap in the arsenite stress- and heat stress-induced ubiquitinomes in both the soluble and pellet fractions (**fig. S1E**). This overlap was evident not only among proteins that showed increased ubiquitination, but also those that showed decreased ubiquitination (**fig. S1F**). Nevertheless, this analysis also confirmed stress-specific patterns in altered ubiquitination. Indeed, more than 400 significant changes in ubiquitination were unique to either arsenite or heat stress (**fig. S1, E and F, tables S2 and S5**).

### Stress-dependent ubiquitination is not reflected by altered abundance

We next sought to determine the extent to which changes in the stress-induced ubiquitinomes reflected changes in total protein levels due to altered rates of synthesis or ubiquitin-dependent degradation. To investigate changes in expression at the protein level, we performed multiplexed TMT MS quantification of the whole proteome from unstressed, heat-stressed, or arsenite-stressed cells in the presence or absence of proteasome inhibition (0.5 μM bortezomib) (**Fig. 3A**). Lysates were digested and labeled by TMT in 11-plex mode, followed by nanoscale LC-high resolution MS/MS. Remarkably, of the 12,586 proteins quantified, the vast majority of proteins showed no statistically significant stress-dependent change in total abundance using 1.5-fold and *P* ≤ 0.05 as a cut-off (**Fig. 3B**, **fig. S2A**, **and table S6**). Nevertheless, we detected several expected changes in total protein levels in response to both stress types, including the accumulation of the well-known stress response transcription factors ATF4 and XBP1, and depletion of BTG1, a regulator of ATF4 (*44*) (**fig. S2B and table S6**). Importantly, no proteins with a stress-dependent increase in ubiquitination showed a significant increase in total protein levels, indicating that increased capture in the TUBE experiments was reflective of increased ubiquitination rather than increased levels of total protein. Additionally, only eight proteins with heat shock-dependent increases in ubiquitination showed a statistically significant decrease of ≥ 1.25-fold in total protein levels in response to stress, of which only three proteins (EPPK1, GCN1, CYP51A1) decreased ≥ 1.5-fold (**Fig. 3B**, **left panel, blue dots, and fig. S2B**). The levels of these three proteins were stabilized during heat shock by the addition of bortezomib, indicating that they are targets of heat stress-dependent proteasomal degradation (**Fig. 3B, right panel, and fig. S2B**). Nonetheless, for nearly all cases with heat shock (**Fig. 3B**) and all cases with arsenite (**fig. S2A**), a stress-dependent increase in ubiquitination did not significantly alter the total cellular pool of a given protein. Notably, treatment with bortezomib during the 1-hour heat shock did not significantly alter ubiquitinated protein enrichment based on TUBE pulldown (**fig. S2C and D**), indicating no substantial degradation of the heat shock ubiquitinome during this time frame. These results suggest that either the observed ubiquitination is non-degradative or that only a minor fraction of each protein type is ubiquitinated in response to stress. Similarly, RNA-seq analysis revealed that most heat shock ubiquitinome genes are not regulated at the level of transcription, with a few exceptions (**Fig. 3C, fig. S2E, table S7**). We further noted that upregulation of many heat shock response genes at the mRNA level (e.g., *HSPA6*, *DNAJB1*, **table S7**) was not yet apparent at the protein level at the 1-hour time point used here (**table S6**), consistent with previous reports that demonstrate a lag time between stress-dependent induction at the transcriptional and translational levels (*45–47*).

**Fig. 3.**
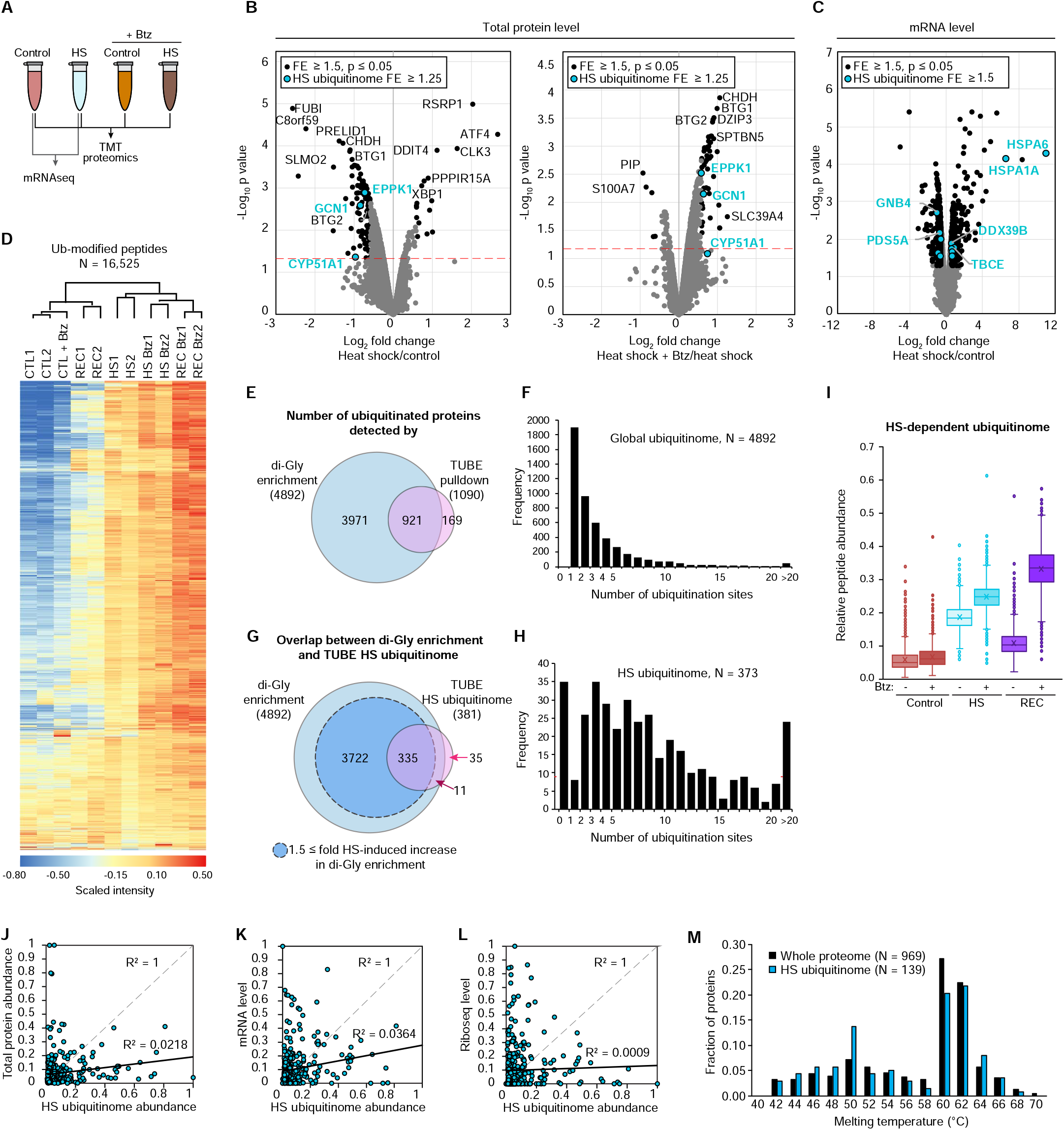
Total proteomic, transcriptomic, and di-GLY analyses reveal additional details of the heat shock ubiquitinome. (**A**) Workflow for total proteome and transcriptome analysis. (**B**) Volcano plots indicating changes in total protein abundance for heat shock versus control samples (left) and heat shock versus heat shock in the presence of the proteasome inhibitor bortezomib (Btz) (right). Statistically significant changes with the indicated parameters are shown for the whole proteome (black dots) and heat shock ubiquitinome (blue dots). Red dashed lines indicates *P* = 0.05. (**C**) Volcano plot indicating statistically significant changes in mRNA levels in response to heat shock. *HSPA1A* and *HSPA6* were the most upregulated genes that were also in the heat shock ubiquitinome (see also fig. S2E); four other representative proteins are indicated. (**D**) Heatmap illustrating global changes to the ubiquitinome during heat shock (HS) and recovery (REC) in the presence or absence of bortezomib (Btz) as determined by di-GLY profiling. Color indicates scaled TMT intensity. (**E**) Venn diagram showing overlap of ubiquitinated proteins detected in di-GLY and TUBE experiments. (**F**) Histogram showing the number of ubiquitination sites per protein detected for all proteins in di-GLY experiments. (**G**) Venn diagram showing overlap of heat shock-dependent increases in ubiquitination as determined by di-GLY and TUBE experiments. (**H**) Histogram showing the number of ubiquitination sites per protein for the heat shock ubiquitinome. (**I**) Box-and-whisker plots showing relative abundance of ubiquitinated peptides for heat shock ubiquitinome proteins. Total peptide abundance (TMT intensity) in all samples was normalized to 1 for each peptide. (**J-L**) Correlation between abundance of heat shock ubiquitinome proteins as determined by TUBE experiments and total protein abundance (J), transcript abundance (K), and ribosome occupancy (L) as determined by Ribo-seq, with R^2^ values displayed. (**M**) Histogram showing melting temperatures as determined in reference (*50*) for all proteins measured (black bars) and measured heat shock ubiquitinome proteins (blue bars).

### Paired di-GLY and TMT quantitative proteomics indicate that heat stress-induced ubiquitination leads to degradation during the recovery phase

Di-GLY proteomics is an alternative approach to identify and quantify the ubiquitin-modified proteome that can validate TUBE-based proteomics and identify specific sites of ubiquitin conjugation (*39, 48*). This approach uses antibody-based affinity to enrich the Lys-ϵ-Gly-Gly (di-GLY) remnant that is generated following trypsin digestion of ubiquitinated proteins, and these peptides are then identified by mass spectrometry. We paired di-GLY proteomics with TMT labeling in the presence or absence of proteasome inhibition to quantify changes in the levels of ubiquitinated peptides in response to a 60-minute heat stress and also following a 2-hour recovery period. Across all samples, we quantified 16,525 unique ubiquitin-modified peptides, corresponding to 4,892 unique proteins (**Fig. 3D and table S8**). We found a high degree of correlation between heat stress-induced ubiquitinomes as determined by di-GLY/TMT proteomics and our earlier TUBE-based proteomics, detecting at least one ubiquitin-modified peptide for the majority of proteins identified by TUBE pulldown (**Fig. 3, E and F and tables S2, S8**). Of the 381 proteins in the heat shock ubiquitinome by TUBE pulldown, 335 had at least one peptide increased by ≥ 1.5-fold in response to heat shock in the di-GLY analysis (**Fig. 3G and tables S4 and S8**). Of the 46 that did not, 35 were not detected at all in the di-GLY experiment, 8 had overall reduced ubiquitinated peptide levels in response to heat shock (HS/CTL = 0.53-0.95), and 3 had modestly increased levels (HS/CTL = 1.29-1.33) (**Fig. 3G and tables S4 and S8**). We note that the amplitudes of the observed changes were often much lower in the di-GLY profiling experiment, likely due to ratio suppression commonly observed in TMT-based quantification (**tables S2 and S8**) (*49*). Interestingly, the heat shock ubiquitinome had much higher numbers of ubiquitination sites per protein (8.6) than the global ubiquitinome (3.4) (**Fig. 3, F and H**), suggesting concerted ubiquitination of a subset of proteins that define the heat shock ubiquitinome. Due to the increased depth of coverage, we identified an additional 3,722 proteins not detected in the TUBE spectral counting for which ubiquitin-modified peptide levels were increased by at least 1.5-fold in response to heat shock (**Fig. 3G**). While the additional heat shock-dependent ubiquitination events detected only by di-GLY profiling represent interesting biology that warrants further investigation, the TUBE-based experiments identified a critical subset of the heat shock ubiquitinome that allowed for comparison between many more sample conditions (e.g., soluble vs pellet, arsenite vs other stresses, multiple cell lines) due to the vastly reduced time and resources required. For this reason, we adjusted the definition of the “heat shock ubiquitinome” as those proteins detected in the TUBE experiments as described above but excluding the 8 proteins showing a heat shock-dependent decrease in the di-GLY experiment (373 proteins, **table S4**).

We next analyzed the di-GLY data to determine the effect of proteasome inhibition on the levels of ubiquitin-modified peptides during heat shock and recovery (**Fig. 3I**). In accordance with our TUBE experiments and whole proteome analysis, we observed that treatment with bortezomib had, at most, a modest effect on abundance of peptides in the heat-shocked samples for most heat shock ubiquitinome proteins. Interestingly, while most of heat shock-dependent ubiquitination recovered to baseline levels after 2 hours in the absence of proteasomal inhibition, the levels remained elevated during recovery in the presence of bortezomib. This result suggests that many proteins ubiquitinated during continued heat stress are likely ultimately targeted for proteasomal degradation upon recovery.

### The heat shock ubiquitinome is not defined by the most abundant, highly translated, or thermo-labile proteins

We next sought to elucidate general features of the heat shock ubiquitinome by analyzing data from the whole-proteome TMT and RNA-seq dataset along with several proteome-wide datasets. We note that the label-free MS approach used in our TUBE enrichment experiments is inherently biased toward identifying abundant proteins. However, proteins detected in the TUBE pulldown experiment spanned four orders of magnitude in TMT intensity, indicating that the heat shock ubiquitinome does include low abundance proteins (**tables S2 and S6**). Furthermore, the spectral count values for the proteins detected in the TUBE pulldown experiment were largely uncorrelated with total protein or mRNA abundance (**Fig. 3, J and K**). This observation, along with the high degree of correlation with the di-GLY profiling, indicates that the quantification of TUBE pulldown samples reflects levels of ubiquitinated protein rather than total protein or transcript levels. Similarly, based on analysis of ribosome profiling data from control HEK293T cells, we found no correlation between the TUBE-based spectral counting quantification of heat shock ubiquitinome proteins and density of ribosome occupancy for the corresponding mRNA (**Fig. 3L**), suggesting that the heat shock ubiquitinome is not composed of DRiPs arising from the most actively translated proteins. Finally, we compared the heat shock ubiquitinome to a previous proteome-wide investigation of protein thermostability (*50*). This published dataset included 139 proteins from the heat shock ubiquitinome, which showed a very similar bimodal distribution of melting temperatures as compared to all 969 proteins analyzed in the study (**Fig. 3M**). Thus, it is unlikely that the heat shock ubiquitinome represents the proteins most likely to misfold as a result of temperature increase. Together these data suggest that the heat shock ubiquitinome is not primarily defined by total protein abundance, translation activity, or thermostability.

### The functional landscape of the heat shock ubiquitinome reflects cellular activities perturbed by stress

To investigate the functional significance and biological implications of the heat shock ubiquitinome, we performed gene ontology (GO) analysis (DAVID (*51*) and PANTHER (*52*)) along with literature curation (**Fig. 4, A-C**). Remarkably, this analysis revealed that the heat shock ubiquitinome is a non-random group of proteins highly enriched in biological pathways that are downregulated or shut down during cellular stress. Furthermore, we detected enrichment of proteins involved in stress granule formation, including translation initiation factors and other components of the translational machinery (**Fig. 4C**). We also noted that the heat shock ubiquitinome includes several classes of nucleocytoplasmic transport proteins (e.g., 11 of the 18 importin α/β superfamily members, components of the nuclear pore complex, and RNA transport proteins), consistent with a recent report that stress disrupts nucleocytoplasmic transport through localization of transport factors into stress granules (*8*) and further linking ubiquitination with downregulated cellular activities and stress granules (**Fig. 4C**). Collectively, these results suggest that in addition to responding to increased PQC demands, ubiquitination plays a role in regulating a variety of biological activities that comprise the cellular stress response, including stress granule assembly. Furthermore, these results suggest that ubiquitination may also contribute to previously underexplored aspects of the heat stress response such as folate and cholesterol metabolism (**Fig. 4C**).

**Fig. 4.**
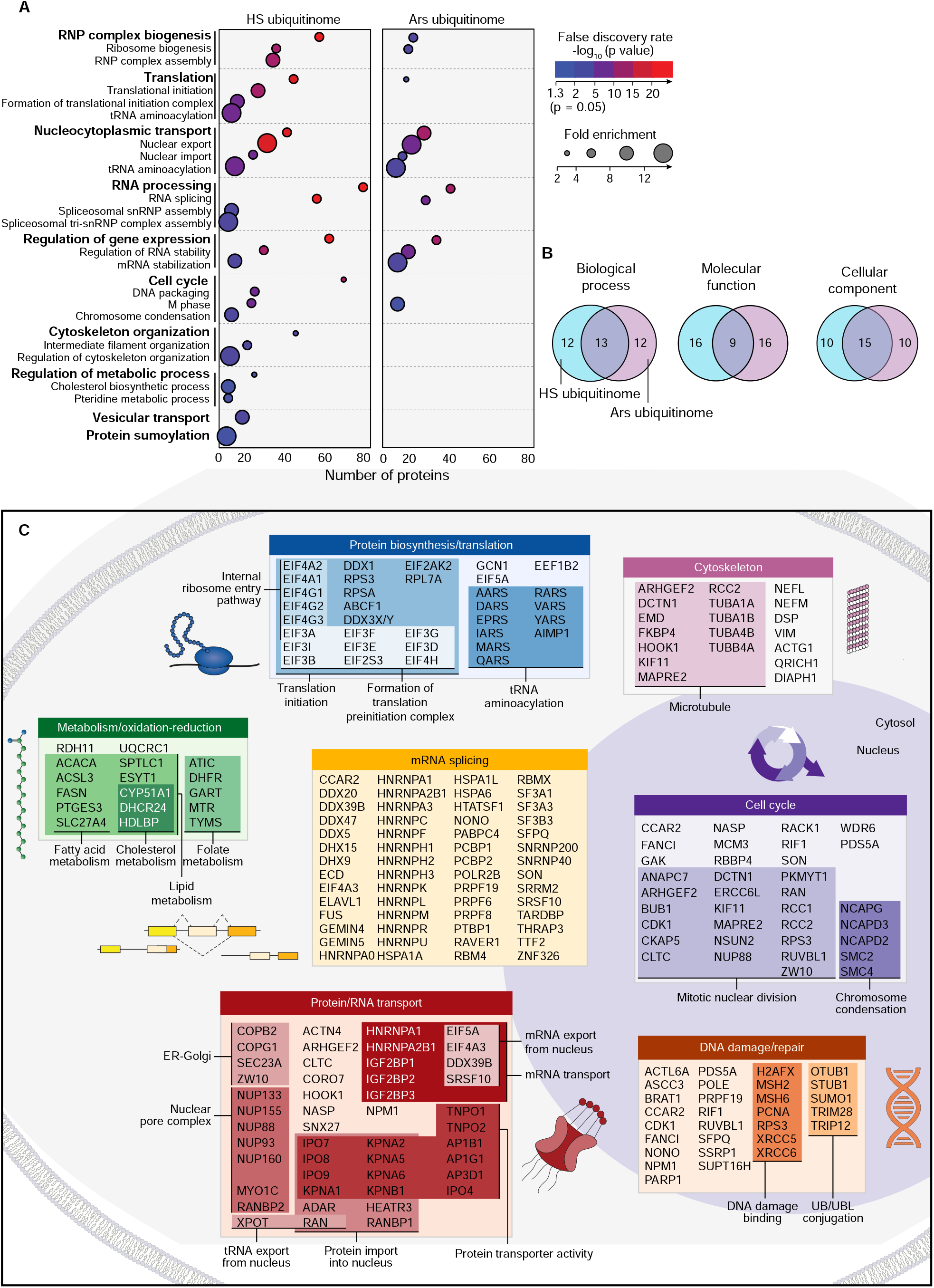
The heat shock ubiquitinome consists of proteins from biological pathways associated with the stress response. (**A**) Dot plot of gene ontology (GO) enrichment showing significantly overrepresented GO biological process terms for heat shock and arsenite ubiquitinomes. Color indicates FDR *P* value and dot size indicates overrepresentation fold enrichment compared to the whole genome. Dots are not shown for terms with no statistically significant (*P* < 0.05) enrichment. (**B**) Overlap of top 25 GO as terms ranked by statistical significance (FDR *P* value) for heat shock and arsenite ubiquitinomes. (**C**) DAVID functional clustering along with literature curation was used to identify biological pathways represented by heat shock ubiquitinome proteins.

Comparisons of the GO analysis of the heat shock- and arsenite-induced ubiquitinomes showed overlap of many common GO terms (e.g., splicing, nucleocytoplasmic transport) and others that were unique to heat shock (e.g., translation, metabolic process) (**Fig. 4, A and B**), suggesting that ubiquitination may play distinct roles in the response to each stress type. Notably, within the GO categories shared by both ubiquitinomes, the heat shock ubiquitinome generally included both a higher number of proteins and higher levels of enrichment over control samples compared to the arsenite ubiquitinome (**Fig. 4A and tables S2, S4 and S5**).

### mRNPs are ubiquitinated *en bloc* and have an altered interactome upon heat shock

Conspicuously, a large fraction of the heat shock ubiquitinome was mRNA-binding proteins (**Fig. 4**), which raised the possibility of *en bloc* ubiquitination of mRNP complexes. To examine this possibility, we first captured mRNPs using magnetic beads conjugated with oligo(dT)25, which binds to the poly(A) tail of mRNAs, and performed immunoblotting to test for the presence of mRNA bound to poly-ubiquitinated proteins (**Fig. 5, A and B**). Results from this assay indicated that poly-ubiquitinated proteins co-purified with polyadenylated mRNA to a much greater extent after heat stress. Poly(A) binding protein (PABP)-mediated 3′ end retrieval is an orthologous method for capturing polyadenylated mRNAs (*53*). When we isolated mRNPs by immunoprecipitating endogenous PABPC1, we similarly observed substantial heat stress-induced poly-ubiquitin conjugates that co-purified with mRNPs (**Fig. 5, A and B**). Importantly, arsenite stress did not lead to a similar level of increased poly-ubiquitin associated with PABPC1, nor did treatment with translation inhibitors cycloheximide, puromycin, or cephaeline in the absence of heat stress, indicating that this phenomenon is specific to heat stress rather than a general response to stress or inhibition of translation (**fig. S3A**).

**Fig. 5.**
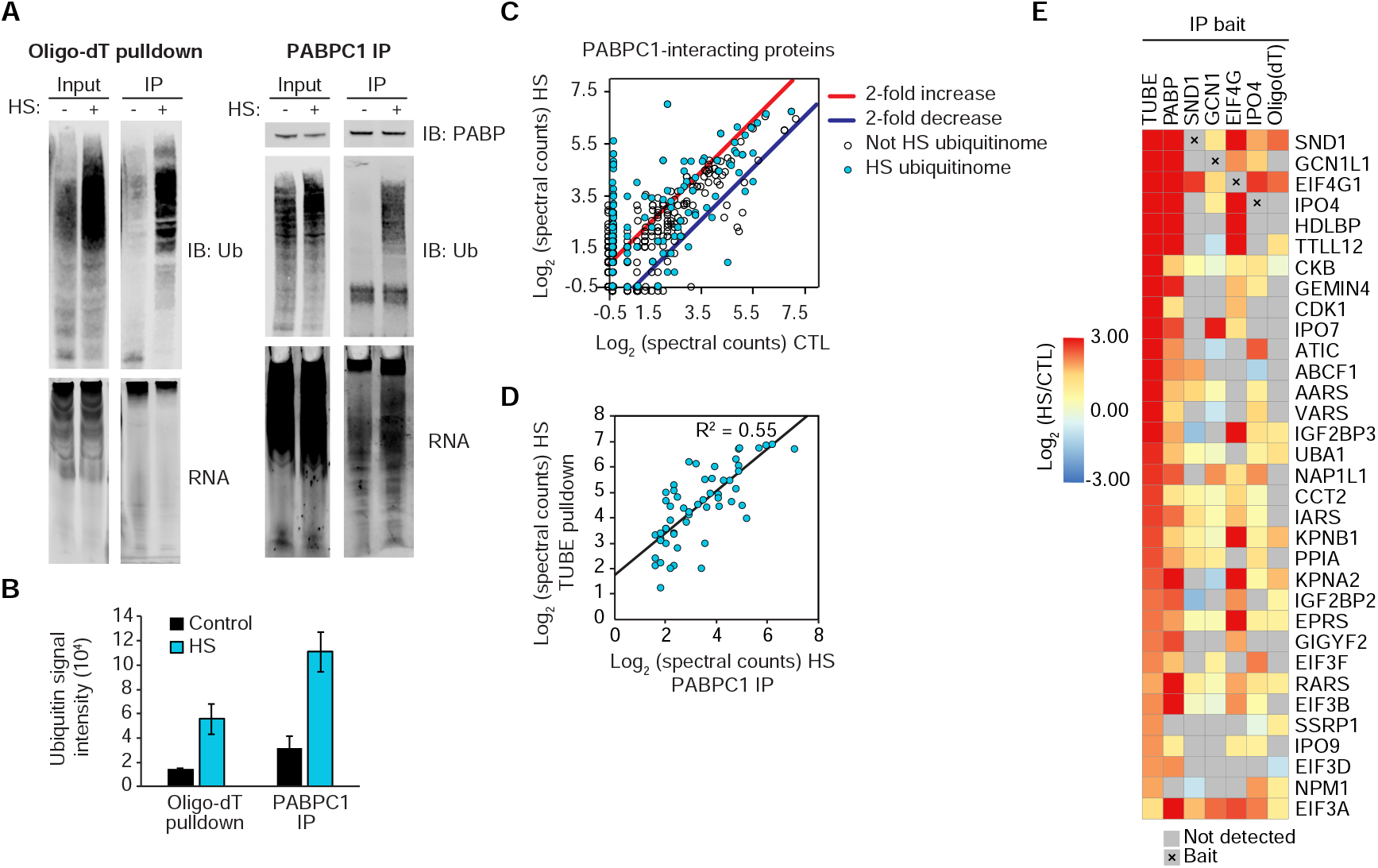
Heat shock induces ubiquitination of mRNP complexes. (**A**) Immunoblots and RNA gels showing polyadenylated mRNA isolated by oligo(dT) resin (left) or PABPC1 immunoprecipitation (right) pulling down poly-ubiquitinated proteins after 60 min heat shock. Isolated mRNA was visualized by SYBR orange RNA stain. (**B**) Quantification of immunoblot analysis for poly-ubiquitin shown in (A). Results represent average of 3 replicate experiments. Error bars indicate s.d. (**C**) Scatter plot showing abundance (log_2_ [spectral counts]) of PABPC1-interacting proteins in heat shock versus control conditions. Results represent averages of 2 replicates. Lines indicate 2-fold increase (red) or decrease (blue) with heat shock; heat shock ubiquitinome proteins are indicated by blue dots. (**D**) Correlation between abundance of heat shock ubiquitinome proteins detected in TUBE pulldown and PABPC1 IP in 60 min heat-shocked samples with indicated R^2^ value. (**E**) Heatmap illustrating changes in abundance in 60 min heat shock versus control samples for proteins detected in TUBE pulldown, endogenous protein IP (PABP, SND1, GCN1, EIF4G, IPO4), or oligo(dT) pulldown.

Direct analysis of the PABPC1 protein-protein interaction network from IP of the endogenous protein revealed substantial remodeling in response to heat stress. Indeed, 105 proteins showed significantly increased interaction with PABPC1 after heat shock, whereas 18 proteins showed significantly decreased interaction (*P* ≤ 0.02, fold enrichment/depletion ≥ 2, detected in both replicates) (**Fig. 5C and table S9**). More than half of the proteins that exhibited a heat stress-dependent increase in co-immunoprecipitation with PABPC1 (59 of 103) were components of the heat shock ubiquitinome (**Fig. 5C)**. Moreover, the relative amount of protein retrieved in PABPC1 immunoprecipitation after heat stress strongly correlated with abundance in the TUBE pulldown experiment (**Fig. 5, D and , E tables S2 and S9**). Consistent results were obtained with 4 additional bait proteins, substantiating the idea of *en bloc* ubiquitination of mRNPs (**Fig. 5E and table S9**). Notably, some of the interactions in the co-IP experiments were observed at low levels in control samples but increased with heat shock (e.g., GCN1 with EIF4G1), whereas others were only detected after heat shock (e.g., TTLL12 with PABPC1 and EIF4G1) (**Fig. 5E and table S9**). One possibility to emerge from this series of experiments is stabilization of PABPC1 complexes through direct, poly-ubiquitin-mediated interactions. However, when we repeated the PABPC1 immunoprecipitation with the addition of the nonspecific deubiquitinating enzyme USP2 that was sufficient to eliminate poly-ubiquitin, we saw no change in the PABPC1 interactome (**fig. S3B**). Altogether, these results suggest *en bloc* ubiquitination of protein components of mRNPs, with some pre-existing interactions that become stabilized upon heat shock as well as some novel heat shock-dependent interactions, reflecting remodeling of the mRNPs.

### Ubiquitination is required for the disassembly of heat shock-induced stress granules

Our initial analysis of the heat shock ubiquitinome highlighted strong enrichment for protein constituents of stress granules. To pursue this observation, we assembled a list of 726 proteins found in stress granules as reported in three independent studies (*25, 26, 28*) (**table S10**). Interestingly, we observed that many more stress granule proteins were represented in the heat shock ubiquitinome than the arsenite ubiquitinome (**Fig. 6, A and B**). For a high confidence stress granule proteome list (those detected in at least 2 of the 3 aforementioned investigations) (**tables S2, S5, and S10**), we observed a clear overall increase in stress granule protein ubiquitination in response to heat shock (**Fig. 6C**). Consistent with a previous report (*32*), this preferential increase in ubiquitination of stress granule proteins was not apparent in response to arsenite stress (**Fig. 6D**). Confirming these results, we observed robust co-localization of poly-ubiquitin with stress granule markers induced by heat shock, but not in those induced by arsenite (**Fig. 6, E and F**).

**Fig. 6.**
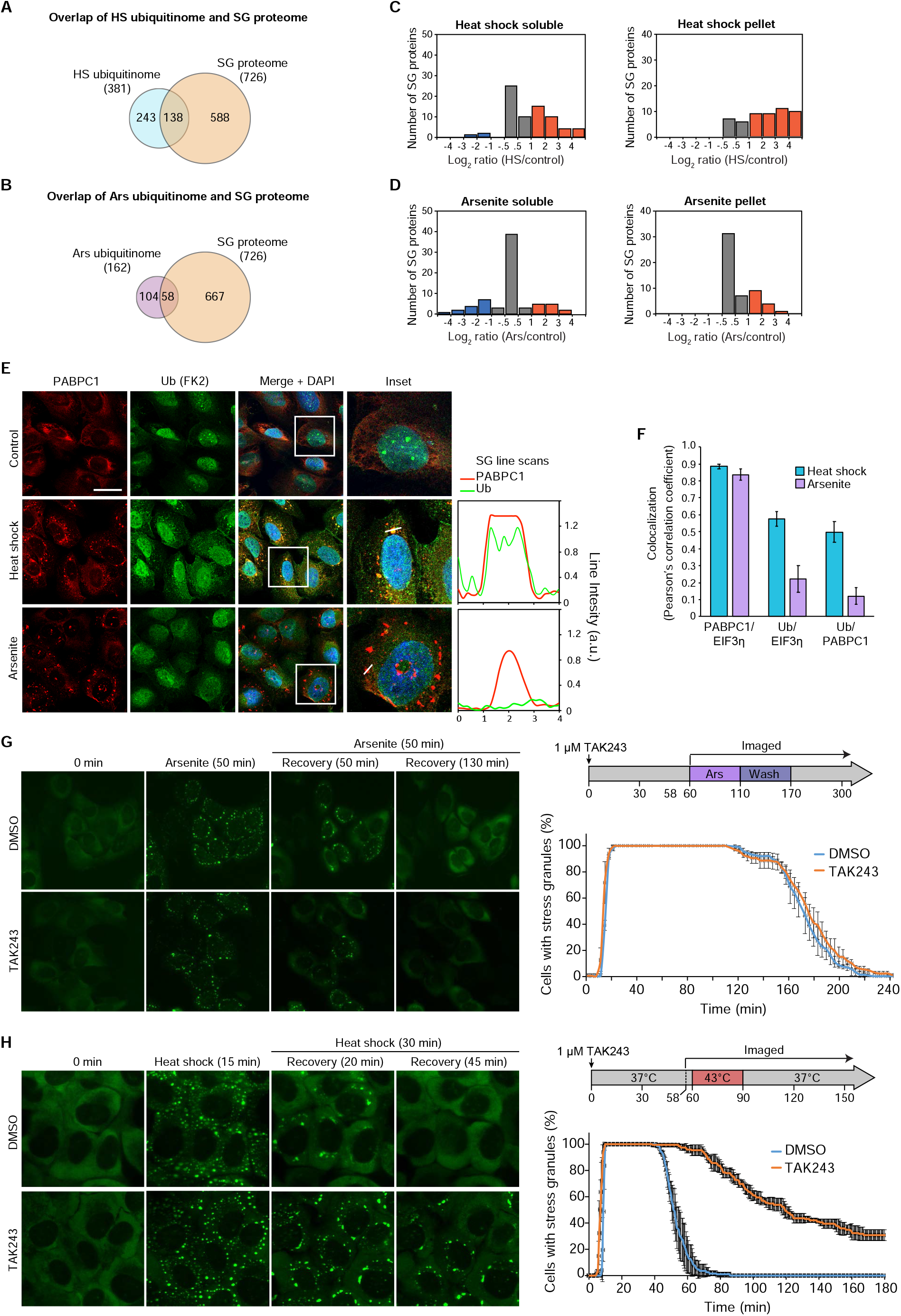
Heat shock-induced stress granules contain ubiquitinated proteins and require active ubiquitination for disassembly. (**A-B**) Overlap of heat shock (A) and arsenite (B) ubiquitinomes with the stress granule proteome. (**C-D**) Histograms showing changes in ubiquitinated protein abundance for high-confidence stress granule proteins in response to 60 min heat shock (C) or 60 min arsenite (D) in soluble and pellet fractions as determined by TUBE experiments (tables S2 and S5). Orange and blue bars indicate > 2-fold increase or decrease respectively. (**E**) U2OS cells were exposed to 90 min of sodium arsenite or heat shock, fixed, and stained for stress granule markers PABPC1 (red), EIF3η (not shown), and poly-ubiquitin (FK2, green). Insets show a magnified image of a stress granule; graphs represent a line scan of signal intensity for PABPC1 and FK2 channels across the indicated stress granule. Scale bar, 50 μm. (**F**) Co-localization between immunofluorescent signal for stress granule markers PABPC1 and EIF3η and poly-ubiquitin as determined from U2OS images. Bar graphs represent average values; error bars represent s.d. of Pearson’s correlation coefficient for stress granules from > 100 cells for each condition. (**G-H**) Live cell imaging of U2OS cells stably expressing GFP-G3BP1 during arsenite (G) or heat (H) stress and recovery in the presence or absence of TAK243. Still images are shown at indicated times (left). Plots show the percentage of cells with ≥ 2 stress granules at the indicated time, with solid lines and error bars representing average values and s.d. for three biological replicates with 30-50 cells each.

To further investigate the relationship between poly-ubiquitin conjugation and stress granule dynamics, we made use of the recently developed E1 ubiquitin activating enzyme (UBA1) inhibitor, TAK243 (*54*). First, we verified that TAK243 treatment led to a dose-dependent and time-dependent depletion of cellular ubiquitin conjugates (**fig. S4**). It was previously reported that elimination of ubiquitination with TAK243 did not influence the assembly or disassembly of arsenite-induced stress granules (*32*). We independently replicated this result using live cell imaging, confirming that ubiquitination does not impact the dynamics of stress granules induced by arsenite (**Fig. 6G and movies S1, S2**). However, given the differences between the ubiquitin response to heat shock and arsenite, we next assessed the role of ubiquitination in regulating the assembly and disassembly of heat shock-induced stress granules. Indeed, considering the prominent ubiquitination of mRNPs we observed following heat stress, we suspected that ubiquitination of mRNPs might be important for stress granule assembly in this setting. To our surprise, elimination of ubiquitination with TAK243 did not prevent stress granule formation in response to heat shock, but rather caused a striking impairment in stress granule disassembly after the removal of stress (**Fig. 6H and movies S3, S4**). We confirmed this observation with immunofluorescence imaging of 10 additional stress granule markers, which showed persistence in stress granules for up to two hours after restoration of normal growth temperature (**fig. S4, B-D**). We also examined how mitigating stress granule assembly impacted the heat shock ubiquitinome. Consistent with the idea that ubiquitination of mRNPs occurs upstream and independent from stress granule formation, we found that blocking stress granule assembly either pharmacologically (i.e., cycloheximide (*55*)) or genetically (i.e., *G3BP1/2* double KO cells (*56*)) had no detectable effect on the heat shock ubiquitinome (**fig. S4, E-G and table S3**). These data indicate that, whereas ubiquitination is not required for heat stress-induced stress granule assembly, it is essential for their rapid disassembly during recovery from stress.

### Reversal of mRNP remodeling following heat shock requires ubiquitination

Given our surprising finding that the disassembly of heat shock-induced stress granules was dependent on ubiquitination, we next investigated the importance of ubiquitination in the reversibility of the heat shock-dependent interactions we observed in the remodeled mRNPs. We added 1 μM TAK243 to the culture media prior to and during heat shock and confirmed elimination of the poly-ubiquitin signal associated with either PABPC1 or oligo(dT) retrieval (**Fig. 7A and fig. S5**). TAK243 treatment did not block the heat shock-dependent remodeling of the mRNP interactome; however, similar to what we observed with stress granules, elimination of ubiquitination prevented the reversal of this stress-dependent remodeling during recovery (**Fig. 7A and fig. S5**). These results indicate that ubiquitination is required not only for disassembly of stress granules, but also the reversibility of mRNP remodeling following restoration of normal growth conditions after heat shock.

**Fig. 7.**
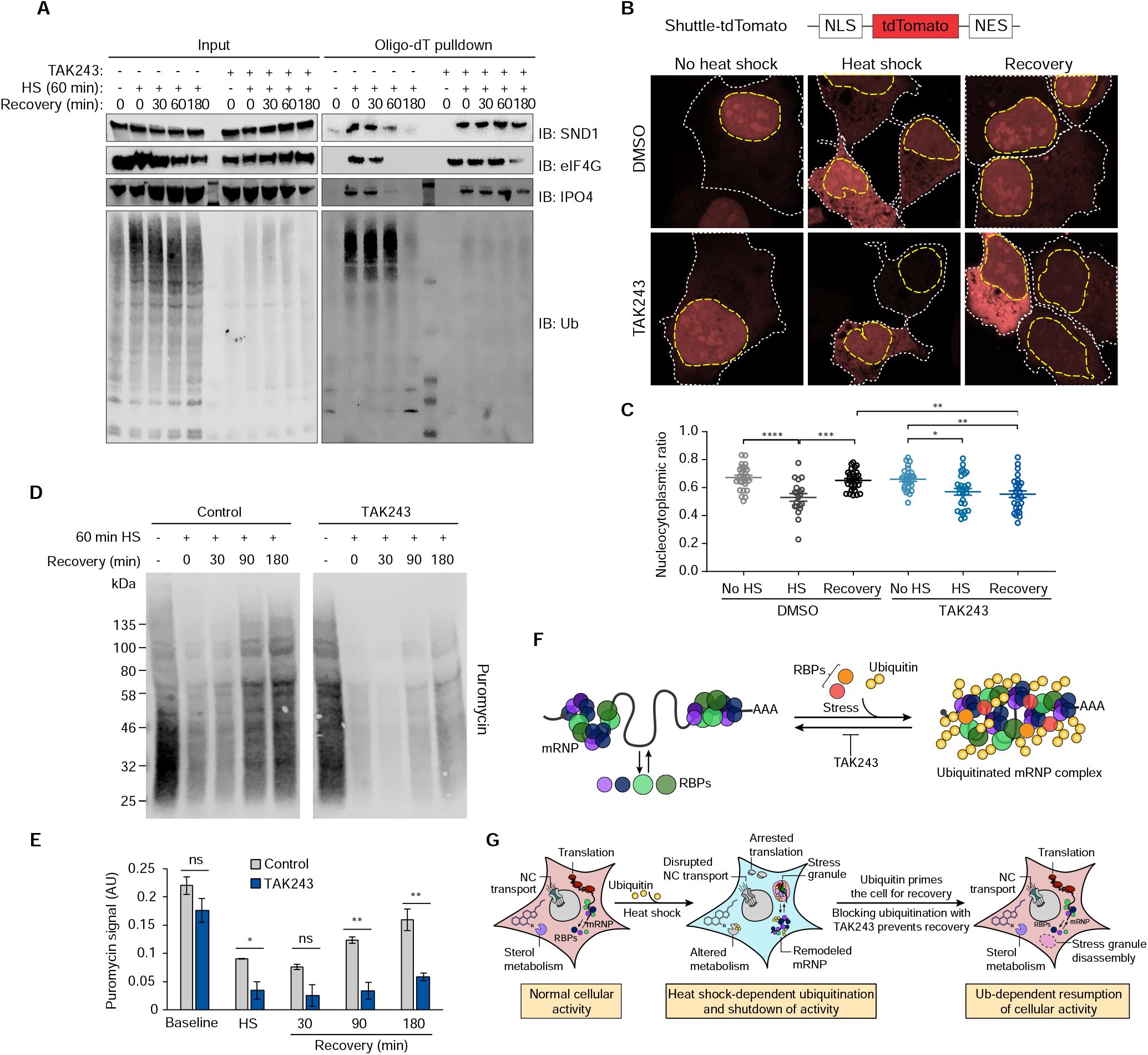
Ubiquitination is required for the recovery of nucleocytoplasmic transport and translation following heat shock. (**A**) Immunoblot showing formation and dissolution of poly-ubiquitinated protein-mRNA complex during heat shock and isolated by oligo(dT) resin. Where indicated, TAK243 was added to cell culture media 30 min prior to heat shock and incubation with the drug was maintained for the duration of the experiment. In drug-treated unstressed samples, cells were incubated with TAK243 for 180 min prior to lysis. (**B**) HEK293T cells expressing nucleocytoplasmic transport reporter NLS-tdTomato-NES were fixed and imaged after no stress, 60 min heat shock, or 60 min heat shock and 120 min recovery. Cells were treated with TAK243 or DMSO for 30 min prior to heat shock or for a total of 120 minutes in non-heat-shocked cells. Nuclear and cytoplasmic boundaries are indicated with dashed yellow and white lines respectively. (**C**) Quantification of nucleocytoplasmic ratio of tdTomato intensity from cell images described in (B) from 20-30 cells for each condition. Error bars indicate s.e.m. \**P* < 0.05, \*\**P* < 0.01, \*\*\**P* < 0.001, \*\*\*\**P* < 0.0001, ANOVA with Tukey’s test. (**D**) Immunoblotting of HEK293T cells treated with puromycin to label nascent transcripts for 30 min prior to heat shock and recovery in the presence or absence of TAK243. Cells were lysed at indicated times and translational activity was analyzed by immunoblotting for puromycin. (**E**) Quantification of immunoblots shown in (D). Average puromycin signal is shown for three replicate experiments. Error bars indicate s.d. n.s., not significant, \**P* < 0.05, \*\**P* < 0.01, student’s t-test. (**F-G**) Proposed model illustrating heat shock-induced ubiquitination associated with (F) mRNP remodeling, and (G) stress granule formation and shutdown of cellular activities, and the requirement for active ubiquitination for reversal of these processes during recovery.

### Ubiquitination is required for recovery of heat shock-induced disruption of nucleocytoplasmic transport

Cellular stresses perturb nucleocytoplasmic transport, resulting in redistribution of many proteins between the nuclear and cytoplasmic compartments (*12*). Heat stress-induced inhibition of nucleocytoplasmic transport, particularly nuclear import, is accompanied by collapse of the Ran gradient (*57*), nuclear retention of importin alpha (*13*), and sequestration of multiple nuclear transport factors in stress granules (*8*). Upon removal of stress, normal nucleocytoplasmic transport is restored and the distribution of proteins between the nuclear and cytoplasmic compartments returns to baseline. To investigate the role of ubiquitination in stress-induced perturbation of nucleocytoplasmic transport, we used a Shuttle-tdTomato nucleocytoplasmic reporter assay in which the fluorescent protein tdTomato was conjugated to both a nuclear localization signal (NLS) and nuclear export signal (NES) that mediate its shuttling between the nucleus and cytoplasm (*8*). Under normal growth conditions, the relative strength of the NLS over the NES led to a preferential localization of fluorescent signal to the nucleus, yielding a steady-state ratio of nuclear to total fluorescence (N/W) of ~0.75 (**Fig. 7, B and C**). Upon heat shock, a defect in nuclear import activity led to a significant redistribution of the reporter from the nucleus to the cytoplasm (N/W ~0.5) (**Fig. 7, B and C**). Under normal growth conditions, inhibition of ubiquitination with TAK243 had no significant impact on the steady-state ratio of nuclear versus cytoplasmic reporter signal, indicating that ubiquitination is not a critical determinant of canonical nuclear import or export. Moreover, following elimination of ubiquitination with TAK243, cells exposed to heat shock still showed a defect in nuclear import activity, indicating that ubiquitination is not required for this inhibition. However, cells treated with TAK243 failed to recover from impaired nucleocytoplasmic shuttling even upon restoration of normal growth conditions for 2 hours. Thus, similar to disassembly of stress granules and reversal of mRNP remodeling, ubiquitination is essential for recovery of nucleocytoplasmic shuttling following heat stress (**Fig. 7, A and B**).

### Ubiquitination is required for the reinitiating translation following heat shock

Protein synthesis is also strongly perturbed by heat stress (*58*) due to broad shutdown of most translation, with the exception of heat shock proteins whose translation is preserved (*59*). Upon removal of stress, normal translation is restored. To investigate the role of ubiquitination in stress-induced perturbation of translation, we used a metabolic labeling assay (*60*). As expected, upon heat shock, we observed a dramatic decrease in translation activity as measured by puromycin incorporation into nascent translation products, followed by a time-dependent recovery after restoration of normal growth conditions (**Fig. 7, D and E**). Whereas TAK243 treatment had no effect on basal translation levels, heat-shocked cells treated with TAK243 showed a profound failure to recover protein synthesis following restoration of normal growth conditions (**Fig. 7, D and E**). Thus, ubiquitination is essential for the recovery of translation during the recovery phrase from heat shock.

Taken together, our results indicate that ubiquitination is required for recovery from multiple heat stress-induced cellular perturbations, including reversal of mRNP remodeling (**Fig. 7F**), stress granule disassembly, and resumption of nucleocytoplasmic transport and protein synthesis (**Fig. 7G**). Furthermore, our results indicate that the role of stress-induced ubiquitination is context-dependent; indeed, several of these observations did not hold true for arsenite stress. Our extensive proteomics datasets describing the heat shock ubiquitinome also suggest that ubiquitination facilitates other aspects of the stress response that have yet to be elucidated. Indeed, in preliminary studies exploring the role of ubiquitination on sterol metabolism, we observed that TAK243 treatment had significant effects on the cholesterol biosynthesis pathway during both heat shock and recovery in a pattern entirely different than that observed for the other cellular activities investigated above (**Fig. 7G and fig. S6**). The data presented here will therefore likely provide a valuable resource for directing future studies.

## Discussion

To survive harsh conditions, eukaryotic cells have evolved adaptive stress response programs that include induction of stress response genes, downregulation of key cellular activities, and the formation of stress granules. Accompanying these stress responses is a global increase in ubiquitination that has been conventionally ascribed to the need for degradation of misfolded or damaged proteins. Here we profiled stress-induced ubiquitination in response to a variety of different types of stress and present evidence that different stressors each result in a distinct pattern of ubiquitination. We delved deeply into heat shock-induced ubiquitination and demonstrate that a substantial fraction of this ubiquitination does not correlate with protein abundance, translation status, or thermostability. Rather, a large fraction of the ubiquitinome corresponds to stress granules and biological pathways that are perturbed by stress. To our surprise, we discovered that ubiquitination was dispensable for characteristic heat shock responses, including downregulation of nucleocytoplasmic transport and translation, and the formation of stress granules. However, in the absence of heat shock-induced ubiquitination there was a profound failure in recovery from stress, revealing an unanticipated function for ubiquitination in mediating the recovery from heat stress (**Fig 7G**).

The adaptive nature of stress-induced ubiquitination has long been recognized. Particularly with proteotoxic insults such as heat shock, stress induces several types of potentially hazardous proteins (e.g., DRiPs, thermolabile misfolded proteins, and other damaged proteins) that must be cleared to prevent the formation of toxic species (*29–31, 40–42*), and this need is largely met by ubiquitination and subsequent proteasomal degradation of the offending species. Prior global proteomic analyses have revealed that a substantial portion of the proteome becomes transiently insoluble in response to heat stress (*40–43*), which suggested a second mode whereby the cell can temporarily reshape the proteome to survive a stressful period. Our results add a third dimension to these concepts, indicating that changes to the ubiquitinome are tailored to specific stressors and, in the case of heat shock stress, ubiquitination serves to prime the cell for recovery. Furthermore, our direct comparisons of the relationship between ubiquitination and heat shock- or arsenite-induced stress granules highlight important differences in context-dependent mechanisms regulating what otherwise appear to be very similar condensates.

Finally, our combined proteome and ubiquitinome datasets represent a vast repository of potentially interesting findings. Our di-GLY-TMT analysis is especially rich, representing one of the deepest ubiquitinomes published to date, including many heat shock-dependent ubiquitination events that were not reflected in our TUBE results. These datasets are likely to provide a rich resource for follow-up investigations of the role of ubiquitination in the stress response.

## Acknowledgments

We thank A. Ordureau (Harvard) and W. Harper (Harvard) for providing the plasmid encoding His-HALO-TUBE and for suggesting protocols for purification and use of the TUBEs; R. Parker (University of Colorado) for helpful discussions; and J. Messing (St. Jude) and J. Temirov (St. Jude) for assistance with microscopy.

## Author contributions

B.A.M. and J.P.T. conceived the project and wrote the paper with editorial contributions from all authors; B.A.M. performed all experiments except nucleocytoplasmic transport assay, which was performed by K.Z., and di-GLY and TMT experiments, which were performed by A.M. under the supervision of J.P. Data was analyzed by B.A.M., A.M., H.J.K., and J.P.T. J.P.T. supervised the project.

## Data and materials availability

All data is available in the main text or the supplementary materials.

## Supplementary Materials

Materials and Methods

Figures S1-S6

Movies S1-S4

Tables S1-S10

References (*1–9*)

## References and Notes

1. H. P. Harding et al., An integrated stress response regulates amino acid metabolism and resistance to oxidative stress. Mol Cell 11, 619–633 (2003).

2. K. Pakos-Zebrucka et al., The integrated stress response. EMBO Rep 17, 1374–1395 (2016).

3. C. O. Brostrom, C. R. Prostko, R. J. Kaufman, M. A. Brostrom, Inhibition of translational initiation by activators of the glucose-regulated stress protein and heat shock protein stress response systems. Role of the interferon-inducible double-stranded RNA-activated eukaryotic initiation factor 2alpha kinase. J Biol Chem 271, 24995–25002 (1996).

4. T. E. Dever et al., Phosphorylation of initiation factor 2 alpha by protein kinase GCN2 mediates gene-specific translational control of GCN4 in yeast. Cell 68, 585–596 (1992).

5. D. Ron, Translational control in the endoplasmic reticulum stress response. J Clin Invest 110, 1383–1388 (2002).

6. R. C. Wek, H. Y. Jiang, T. G. Anthony, Coping with stress: eIF2 kinases and translational control. Biochem Soc Trans 34, 7–11 (2006).

7. P. D. Lu, H. P. Harding, D. Ron, Translation reinitiation at alternative open reading frames regulates gene expression in an integrated stress response. J Cell Biol 167, 27–33 (2004).

8. K. Zhang et al., Stress Granule Assembly Disrupts Nucleocytoplasmic Transport. Cell 173, 958–971 e917 (2018).

9. M. Furuta et al., Heat-shock induced nuclear retention and recycling inhibition of importin alpha. Genes Cells 9, 429–441 (2004).

10. J. B. Kelley, B. M. Paschal, Hyperosmotic stress signaling to the nucleus disrupts the Ran gradient and the production of RanGTP. Mol Biol Cell 18, 4365–4376 (2007).

11. M. Kodiha, A. Chu, N. Matusiewicz, U. Stochaj, Multiple mechanisms promote the inhibition of classical nuclear import upon exposure to severe oxidative stress. Cell Death Differ 11, 862–874 (2004).

12. S. Kose, N. Imamoto, Nucleocytoplasmic transport under stress conditions and its role in HSP70 chaperone systems. Biochim Biophys Acta 1840, 2953–2960 (2014).

13. Y. Miyamoto et al., Cellular stresses induce the nuclear accumulation of importin alpha and cause a conventional nuclear import block. J Cell Biol 165, 617–623 (2004).

14. R. Shalgi, J. A. Hurt, S. Lindquist, C. B. Burge, Widespread inhibition of posttranscriptional splicing shapes the cellular transcriptome following heat shock. Cell Rep 7, 1362–1370 (2014).

15. K. Yamamoto et al., Control of the heat stress-induced alternative splicing of a subset of genes by hnRNP K. Genes Cells 21, 1006–1014 (2016).

16. A. K. Pearce, T. C. Humphrey, Integrating stress-response and cell-cycle checkpoint pathways. Trends Cell Biol 11, 426–433 (2001).

17. E. Radmaneshfar et al., From START to FINISH: the influence of osmotic stress on the cell cycle. PLoS One 8, e68067 (2013).

18. N. M. Kuhl, L. Rensing, Heat shock effects on cell cycle progression. Cell Mol Life Sci 57, 450–463 (2000).

19. V. Balagopal, R. Parker, Polysomes, P bodies and stress granules: states and fates of eukaryotic mRNAs. Curr Opin Cell Biol 21, 403–408 (2009).

20. C. Jousse et al., Inhibition of a constitutive translation initiation factor 2alpha phosphatase, CReP, promotes survival of stressed cells. J Cell Biol 163, 767–775 (2003).

21. I. Novoa, H. Zeng, H. P. Harding, D. Ron, Feedback inhibition of the unfolded protein response by GADD34-mediated dephosphorylation of eIF2alpha. J Cell Biol 153, 1011–1022 (2001).

22. J. R. Wheeler, T. Matheny, S. Jain, R. Abrisch, R. Parker, Distinct stages in stress granule assembly and disassembly. Elife 5, (2016).

23. X. Xie et al., Deubiquitylases USP5 and USP13 are recruited to and regulate heat-induced stress granules through their deubiquitylating activities. J Cell Sci 131, (2018).

24. S. Kwon, Y. Zhang, P. Matthias, The deacetylase HDAC6 is a novel critical component of stress granules involved in the stress response. Genes Dev 21, 3381–3394 (2007).

25. S. Markmiller et al., Context-Dependent and Disease-Specific Diversity in Protein Interactions within Stress Granules. Cell 172, 590–604 e513 (2018).

26. J. Y. Youn et al., High-Density Proximity Mapping Reveals the Subcellular Organization of mRNA-Associated Granules and Bodies. Mol Cell 69, 517–532 e511 (2018).

27. N. Kedersha et al., G3BP-Caprin1-USP10 complexes mediate stress granule condensation and associate with 40S subunits. J Cell Biol 212, 845–860 (2016).

28. S. Jain et al., ATPase-Modulated Stress Granules Contain a Diverse Proteome and Substructure. Cell 164, 487–498 (2016).

29. A. Turakhiya et al., ZFAND1 Recruits p97 and the 26S Proteasome to Promote the Clearance of Arsenite-Induced Stress Granules. Mol Cell 70, 906–919 e907 (2018).

30. S. Alberti, D. Mateju, L. Mediani, S. Carra, Granulostasis: Protein Quality Control of RNP Granules. Front Mol Neurosci 10, 84 (2017).

31. M. Ganassi et al., A Surveillance Function of the HSPB8-BAG3-HSP70 Chaperone Complex Ensures Stress Granule Integrity and Dynamism. Mol Cell 63, 796–810 (2016).

32. S. Markmiller et al., Active Protein Neddylation or Ubiquitylation Is Dispensable for Stress Granule Dynamics. Cell Rep 27, 1356–1363 e1353 (2019).

33. P. Beaudette, O. Popp, G. Dittmar, Proteomic techniques to probe the ubiquitin landscape. Proteomics 16, 273–287 (2016).

34. A. Ordureau, C. Munch, J. W. Harper, Quantifying ubiquitin signaling. Mol Cell 58, 660–676 (2015).

35. F. Lopitz-Otsoa et al., Integrative analysis of the ubiquitin proteome isolated using Tandem Ubiquitin Binding Entities (TUBEs). J Proteomics 75, 2998–3014 (2012).

36. A. Fulzele, E. J. Bennett, Ubiquitin diGLY Proteomics as an Approach to Identify and Quantify the Ubiquitin-Modified Proteome. Methods Mol Biol 1844, 363–384 (2018).

37. N. N. Fang et al., Rsp5/Nedd4 is the main ubiquitin ligase that targets cytosolic misfolded proteins following heat stress. Nat Cell Biol 16, 1227–1237 (2014).

38. N. N. Fang, M. Zhu, A. Rose, K. P. Wu, T. Mayor, Deubiquitinase activity is required for the proteasomal degradation of misfolded cytosolic proteins upon heat-stress. Nat Commun 7, 12907 (2016).

39. C. M. Rose et al., Highly Multiplexed Quantitative Mass Spectrometry Analysis of Ubiquitylomes. Cell Syst 3, 395–403 e394 (2016).

40. G. Xu et al., Identification of proteins sensitive to thermal stress in human neuroblastoma and glioma cell lines. PLoS One 7, e49021 (2012).

41. E. W. Wallace et al., Reversible, Specific, Active Aggregates of Endogenous Proteins Assemble upon Heat Stress. Cell 162, 1286–1298 (2015).

42. X. Sui et al., Widespread remodeling of proteome solubility in response to different protein homeostasis stresses. Proc Natl Acad Sci U S A 117, 2422–2431 (2020).

43. A. H. Ng, N. N. Fang, S. A. Comyn, J. Gsponer, T. Mayor, System-wide analysis reveals intrinsically disordered proteins are prone to ubiquitylation after misfolding stress. Mol Cell Proteomics 12, 2456–2467 (2013).

44. L. Yuniati et al., Tumor suppressor BTG1 promotes PRMT1-mediated ATF4 function in response to cellular stress. Oncotarget 7, 3128–3143 (2016).

45. K. R. Diller, Stress protein expression kinetics. Annu Rev Biomed Eng 8, 403–424 (2006).

46. J. R. Subjeck, J. J. Sciandra, R. J. Johnson, Heat shock proteins and thermotolerance; a comparison of induction kinetics. Br J Radiol 55, 579–584 (1982).

47. Y. Zhang et al., The role of heat shock factors in stress-induced transcription. Methods Mol Biol 787, 21–32 (2011).

48. G. Xu, J. S. Paige, S. R. Jaffrey, Global analysis of lysine ubiquitination by ubiquitin remnant immunoaffinity profiling. Nat Biotechnol 28, 868–873 (2010).

49. N. Rauniyar, J. R. Yates, 3rd, Isobaric labeling-based relative quantification in shotgun proteomics. J Proteome Res 13, 5293–5309 (2014).

50. P. Leuenberger et al., Cell-wide analysis of protein thermal unfolding reveals determinants of thermostability. Science 355, (2017).

51. W. Huang da, B. T. Sherman, R. A. Lempicki, Systematic and integrative analysis of large gene lists using DAVID bioinformatics resources. Nat Protoc 4, 44–57 (2009).

52. H. Mi et al., PANTHER version 11: expanded annotation data from Gene Ontology and Reactome pathways, and data analysis tool enhancements. Nucleic Acids Res 45, D183–D189 (2017).

53. H. W. Hwang et al., PAPERCLIP Identifies MicroRNA Targets and a Role of CstF64/64tau in Promoting Non-canonical poly(A) Site Usage. Cell Rep 15, 423–435 (2016).

54. M. L. Hyer et al., A small-molecule inhibitor of the ubiquitin activating enzyme for cancer treatment. Nat Med 24, 186–193 (2018).

55. N. Kedersha et al., Dynamic shuttling of TIA-1 accompanies the recruitment of mRNA to mammalian stress granules. J Cell Biol 151, 1257–1268 (2000).

56. P. Yang et al., G3BP1 is a tunable switch that triggers phase separation to assemble stress granules. Cell, (In Press).

57. U. Stochaj, R. Rassadi, J. Chiu, Stress-mediated inhibition of the classical nuclear protein import pathway and nuclear accumulation of the small GTPase Gsp1p. FASEB J 14, 2130–2132 (2000).

58. R. Panniers, Translational control during heat shock. Biochimie 76, 737–747 (1994).

59. R. Panniers, E. C. Henshaw, Mechanism of inhibition of polypeptide chain initiation in heat-shocked Ehrlich ascites tumour cells. Eur J Biochem 140, 209–214 (1984).

60. E. K. Schmidt, G. Clavarino, M. Ceppi, P. Pierre, SUnSET, a nonradioactive method to monitor protein synthesis. Nat Methods 6, 275–277 (2009).

