## Supplemental Methods and Figures for "Ubiquitination is essential for recovery of cellular activities following heat shock"

**This PDF file includes:**

Materials and Methods
Figs. S1 to S6
Caption for Movies S1 to S4
Captions for Tables S1 to S10

**Other Supplementary Materials for this manuscript include the following:**

Movies S1 to S4
Tables S1 to S10 (.xlsx)

### Materials and Methods

Cell culture and stress treatment

HEK293T and U2OS cells were obtained from ATCC. U2OS cells stably expressing G3BP1-GFP have been previously described (*1*). Cells were cultured in Dulbecco's modified Eagle's medium (HyClone) supplemented with 10% fetal bovine serum (Hyclone SH30071.03 and SH30396.03) and maintained at 37°C in a humified incubator with 95% air and 5% $CO_2$. Cells were authenticated by short tandem repeat (STR) profiling. For heat shock treatment, cell culture media was replaced with fresh media pre-warmed to 42°C and placed in a 42°C humidified incubator with 5% $CO_2$ for the indicated time. In recovery experiments, the cells were returned to the 37°C incubator for the indicated time. For matched control-treated cells, media was replaced with fresh media warmed to 37°C and maintained at 37°C for the indicated time prior to harvest. When indicated, 0.5 µM bortezomib was included in the fresh media added to the cells at the start of the experiment. For other stresses, media was replaced with fresh media containing 0.5 mM sodium arsenite, 0.4 M sorbitol, or 1 µM bortezomib for the indicated time. For UV stress, media was removed and replaced with 2 ml PBS and 40 $J/m^2$ UV radiation was delivered using a Stratalinker UV 2400 (Stratagene). Following UV irradiation, PBS was removed, fresh media was added back, and cells were placed in the 37°C incubator for 1 h prior to harvest. When indicated, 1 µM TAK243 was added directly to cell culture media 30 min prior to stress treatment and maintained in the media until lysis. In TAK243-treated non-stressed control samples, TAK243 was added at least 90 min prior to cell lysis as indicated. In puromycin incorporation SUnSET assay, puromycin was added to cell culture media at 1 µg/ml for 15 min prior to cell lysis to label nascent transcripts.

Preparation of cell lysates

Cell pellets were collected by centrifugation at 200 x g followed by removal of media by aspiration, and the cell pellet was washed twice with PBS. Cell pellets were then lysed with urea lysis buffer (30 mM Tris-HCl pH 8.5, 7 M urea, 2 M thiourea, 4% CHAPS), TUBE lysis buffer (50 mM Tris-HCl, 150 mM NaCl, 1% NP-40, 1 mM EDTA, 1 mM EGTA), or IP lysis buffer (20 mM Tris-HCl, 150 mM NaCl, 0.5 % Triton X100) as indicated. All lysis buffers were supplemented with 50 mM iodoacetamide, 50 µM PR-619, 0.5 µM bortezomib, and protease and phosphatase inhibitor mini tablets (Pierce). Lysates were centrifuged for 10 min at 4°C at 20,000 x g and the supernatant fraction was collected as the "soluble fraction" for TUBE or IP lysis buffers or "whole cell lysate" for urea lysis buffer. To collect pellet fractions, the insoluble pellet from TUBE lysis buffer was washed twice with ice-cold PBS and then incubated with 100 µl urea lysis buffer for 10 min with intermittent vortexing prior to a second centrifugation for 10 min at 4°C at 20,000 x g.

TUBE protein purification and conjugation to HaloLink resin

A construct with 4x UBA domains from UBQLN2 (TUBE) cloned into pFN18a-His-HALO was obtained from Wade Harper (Harvard Medical School). The His-Halo-TUBE protein was expressed and purified from *E. coli* Rosetta 2(DE3) cells (Millipore). *E. coli* were grown to $OD_{600}$ of 0.7 and induced with 1 mM IPTG at 25°C overnight. Pelleted cells were resuspended in lysis buffer (50 mM Tris-HCl, pH 7.5, 1% Triton X, 150 mM NaCl, 0.1% (vol/vol) 2-mercaptoethanol, 1 mM benzamidine, and 0.2 mM phenylmethylsulfonyl fluoride (PMSF)) and lysed by sonication. Lysates were clarified by ultracentrifugation at 30,000 x g at 4°C for 1 h. Supernatants were applied to packed Ni columns with 10 ml Ni-NTA Agarose (Thermo Fisher

Scientific) pre-equilibrated with lysis buffer at 4°C. Protein was then eluted with 250 mM imidazole in lysis buffer, dialyzed overnight at 4°C in storage buffer (50 mM Tris-HCl, pH 7.5, 0.15 M NaCl, 10% glycerol, 0.1% 2-mercaptoethanol, 0.1% Triton X-100, 0.2 mM PMSF, 1 mM benzamidine) and stored at -80°C. To conjugate TUBEs to resin, 5 mg of previously purified 6His-Halo-TUBE per ml of HaloLink resin (Promega) was incubated overnight at 4°C with constant mixing. The resin was then washed 5 times with 10 bed volumes storage buffer plus 1 M NaCl and then once with storage buffer and stored at 4°C.

TUBE pulldown

For the soluble fraction, 90% of the supernatant from cells of one 10-cm dish per sample lysed in TUBE lysis buffer was incubated with 50 µl packed volume His-Halo-TUBE beads overnight at 4°C. For the pellet fraction, the pellet from one 15-cm dish in TUBE lysis buffer was resuspended in 100 µl urea lysis buffer as described above. Following a second centrifugation step to remove the urea-insoluble material, the supernatant was diluted 10-fold in TUBE lysis buffer and incubated overnight at 4°C with 50 µl His-Halo-TUBE beads. After the incubation/binding step, a small amount of the supernatant was collected as "unbound sample" for analysis by Western blot. Beads with captured proteins were then washed three times with 1 ml of lysis buffer containing 500 mM NaCl and then once with 10 mM Tris-HCl, pH 8. Captured proteins were then released from the beads by denaturation in 1x Laemmli sample buffer with reducing agent and heating at 75°C for 5 min. Western blot analysis was performed immediately after elution and the remaining sample was stored at -80°C for analysis by mass spectrometry (MS).

Immunoprecipitation and Oligo(dT) pulldown

Cells were lysed in IP lysis buffer and lysates were incubated overnight at 4°C with Protein G Dynabeads (Invitrogen) conjugated to antibodies against endogenous proteins according to the manufacturer's protocol. For Oligo(dT) pulldown, lysate was incubated with 50 µl settled Oligo d(T)25 magnetic bead resin (New England Biolabs) overnight at 4°C. Following overnight incubation, beads were washed 3x with PBS and proteins were eluted in 1x LDS sample buffer with reducing agent and stored at -20°C for analysis by immunoblot and MS.

Western blotting

1X NuPAGE LDS sample buffer and NuPAGE sample reducing agent (Thermo Fisher Scientific, NP0008 and NP0004) were added to cell lysate, TUBE eluate, or IP eluate, and samples were heated at 75°C for 5 min (samples with urea lysis buffer were not heated). Samples were separated in 4-12% NuPAGE Bis-Tris or 3-8% NuPAGE Tris-Acetate gels (Invitrogen) and transferred to nitrocellulose membranes using an iBlot 2 transfer device (Thermo Fisher Scientific). Membranes were blocked with Odyssey blocking buffer (LI-COR) and then incubated with primary antibodies at 4°C overnight. Primary antibodies used in this study are ACTIN (Santa Cruz Biotechnology, sc-1616); ATF4 (Thermo Fisher Scientific, PA5-17989); CYP51A1 (Proteintech, Cat#13431-1-AP); EIF4G1, (Santa Cruz Biotechnology, sc-11373); GAPDH (Santa Cruz Biotechnology, sc-32233); GCN1 (Abcam, ab86139); IPO4 (Sigma Aldrich, SAB4200167); KPNA2 (Abcam, ab70160), PABPC1 (Abcam, ab21060); PD41-ubiquitin (Santa Cruz Biotechnology, sc-8017); Puromycin (EMD Millipore, MABE343); SND1 (Abcam, ab65078); TTLL12 (Thermo Fisher Scientific, MA5-25018); XRCC5 (Thermo Fisher Scientific, MA5-15873). Membranes were washed 3 times with PBST (0.1% Tween) and further incubated with dye-labeled or HRP-conjugated secondary antibodies. Membranes were visualized with an Odyssey Fc imaging system (LI-COR).

Immunofluorescence and microscopy

For fixed cell imaging, cells were grown in 4-well or 8-well chamber slides (Millipore). Cells were fixed with 4% paraformaldehyde (Electron Microscopy Science) in PBS for 10 min, permeabilized with 0.2% Triton X-100 in PBS for 10 min, and then blocked with 1% BSA for 1 h, with all steps performed at room temperature. Samples were further incubated with primary antibodies in blocking buffer overnight at 4°C, then washed three times with PBST (0.1% Tween) and incubated with secondary antibody for 1 h at room temperature. Primary antibodies used in this study are CYP51A1 (Proteintech, Cat#13431-1-AP); EIF3η (Santa Cruz Biotechnology, sc-16377); EIF4G1, (Santa Cruz Biotechnology, sc-11373); EPRS (Abcam, ab31531); FK2-ubiquitin (Enzo Life Sciences, BML-PW8810-0500); G3BP1 (BD Biosciences, Cat#611127); GCN1 (Abcam, ab86139); HNRNPA2B1 (Santa Cruz Biotechnology, sc-32316); IPO4 (Sigma Aldrich, SAB4200167); KPNA2 (Abcam, ab70160); PABPC1 (Abcam, ab21060); SFPQ (Abcam, ab38148); SND1 (Abcam, ab65078); XRCC5 (Thermo Fisher Scientific, MA5-15873). Host-specific Alexa Fluor 488/555/647 (Thermo Fisher) secondary antibodies were used for visualization. For microscopic imaging, slides were mounted with ProLong Gold Antifade reagent with DAPI (Invitrogen). Images were captured using a widefield or laser scanning confocal microscope (Leica) with a 63x 1.4 NA oil objective. For nucleocytoplasmic shuttling assays, HEK293T cells were transfected with pLenti-NLS-tdTomato-NES for at least 24 h prior to treatment and fixation and treated with TAK243 or DMSO for 30 min prior to heat shock. Following treatment, cells were fixed, stained with DAPI, and tdTomato localization was measured in the nucleus and cytoplasm in ImageJ/Fiji. For measuring ubiquitin colocalization into stress granules, U2OS cells were co-stained with antibodies for poly-ubiquitin (FK2), PABPC1, and EIF3η. Line scan intensity profiles were generated in ImageJ/Fiji by measuring intensity along a cross-section of a well-defined stress granule. For quantification of ubiquitin colocalization to stress granules, Ilastik (*2*) was used to identify and segment stress granules from images of at least 100 cells. Then, segmented images were imported into CellProfiler (*3*) along with original images to measure colocalization by determining Pearson's correlation coefficient between protein signals within stress granules in the original images.

Live-cell imaging

Live-cell imaging was performed using U2OS cells stably expressing G3BP1-GFP. For monitoring stress granules during heat shock, cells growing on glass-bottom 35-mm dishes (Corning) were treated with DMSO or TAK243 (1 μM) for 30 min prior to imaging. Imaging was performed using Opterra microscopy (BRUKER) with a cage for environmental control (Oko labs) maintaining 37C and 5% $CO_2$. Heat shock was accomplished with an objective heater (Bioptechs). Using Prairie View software with perfect focus engaged to correct for z-drift, multipoint images were taken every 30 s with the 488-nm laser at 30% power using a 60x 1.42NA objective. Two minutes into imaging, the objective temperature was raised to 43°C. At 30 min after heat shock, the temperature was lowered back to 37°C to alleviate the stress, and cells were imaged until granules disappeared or after 2-3 h had elapsed.

For monitoring arsenite-induced stress granules, cells were grown on 96-well assay plates (Corning 3904) and were treated with DMSO or TAK243 (1 μM) for 30 min prior to imaging. Imaging was performed on a Cytation 5 imaging reader (BioTek) with a 20X Olympus Plan

Fluorite Phase Objective 0.45NA (1320517), using a GFP LED (1225001) and Filter Cube (1225101), along with a Laser Autofocus Cube (1225010) for z-drift correction. Environmental control was maintained within the Cytation at 37°C and 5% $CO_2$. For liquid handling a Biospa 8 (Biotek) and Multiflo FX (Biotek) were attatched to the Cytation 5, with a Cassette 5uL molded tip assembly (Biotek 7170011) present on the Multiflo FX to dispense liquid. Cells were imaged once, followed by arsenite being dispensed at a final concentration of 0.5 mM. Following arsenite addition, cells were imaged for 45 min before being washed out 3 times with media not containing arsenite. Following this the cells were imaged for an additional 45 min, before being washed out 2 times more and returned to Cytation for imaging until granules disappeared. Throughout the dispensing and washing steps, DMSO or TAK243 was maintained at 1 μM in the wells.

**<u>Proteomic profiling by spectral counting</u>**

Mass spectrometric (MS) analysis was performed according to an optimized platform as previously reported (*4*). Eluates from the TUBE pulldown or IP experiments were run on an SDS gel and visualized by Coomassie staining (Thermo Fisher Scientific). The whole gel lanes were excised into gel bands. The proteins in the gel bands were reduced by dithiothreitol (DTT), alkylated by iodoacetamide, and proteolyzed with trypsin at an enzyme-to-substrate ratio of 1:50 (w/w). The digested peptides were then extracted from the gel bands, dried in a speed vacuum, and reconstituted in a loading buffer of 5% formic acid (FA) plus 0.1% trifluoroacetic acid (TFA). The peptides were fractionated on a nanoscale capillary reverse phase C18 column using a 90-min gradient of 7%-35% buffer B (70% acetonitrile (ACN) plus 0.2% FA, with buffer A containing 0.2% FA) at a flow rate of 0.30 µL/min delivered by an HPLC system (Thermo Scientific EasynLC 1000), and detected by an in-line mass spectrometer (Thermo Scientific LTQ Orbitrap Elite) operating in a data-dependent "High-Low" rapid scanning mode, with a survey scan in the Orbitrap (60,000 high resolution, scan range 400–1600 m/z, $1 \times 10^6$ automatic gain control target, ~50 ms maximal ion time), followed by sequential isolation of top 20 abundant ions for MS/MS analysis in the linear ion trap of low resolution (fragmentation by collision-activated dissociation with 35 normalized collision energy, $1 \times 10^5$ automatic gain control (AGC) target, ~100 ms maximal ion time, 2.0 *m/z* isolation window, and 20 s dynamic exclusion). MS/MS spectra were searched against UniProt human protein database using SEQUEST algorithm in our JUMP software package (*5*) (https://github.com/JUMPSuite/jumpsuite.github.io). Search parameters included mass tolerance of 30 ppm for precursor ions and 0.2 Da for MS/MS ions, full trypticity with maximal two missed cleavages, maximal three dynamic modification sites per peptide, carbamidomethylation of Cys (+57.02146 Da) as static modification and Met oxidation (+15.99492 Da) as dynamic modification. All matched MS/MS spectra were filtered by mass accuracy and matching scores to reduce protein false discovery rate (FDR) to about 1%, based on the target-decoy strategy using a reversed database (*6*).

**<u>Proteomic profiling of whole proteome by TMT labeling</u>**

Proteomic profiling of whole proteome was carried out with a previously reported protocol (*7*). Briefly, for each sample, HEK293T cells from one 10-cm dish were harvested, washed with ice-cold PBS and collected by centrifugation. Cells were lysed in a buffer (50 mM HEPES, pH 8.5, 8 M urea, and 0.5% sodium deoxycholate). Protein concentration was estimated by Coomassie staining of a gel where samples were run for a short distance, using bovine serum albumin (BSA) as a standard. About 100 µg of protein per sample was digested with LysC (Wako) at an enzyme-to-substrate ratio of 1:100 (w/w) for 2 h at room temperature, followed by Cys reduction (1 mM DTT for 30 min), alkylation (10 mM iodoacetamide for 30 min) and quenching (30 mM DTT for 30 min) at room temperature. The samples were diluted to a final concentration of 2 M urea with 50 mM HEPES (pH 8.5), and further digested with trypsin (Promega) at an enzyme-to-substrate ratio of 1:50 (w/w) for at least 3 h at room temperature. Finally, the digestion was terminated and acidified by adding TFA to 1%, centrifuged at 21,000 x g for 10 min at 4°C to remove any insoluble material. The supernatant were desalted on C18 SepPak solid-phase extraction cartridges (SPE) (Waters), dried, resuspended in 50 mM HEPES (pH 8.5), and differentially labeled with 11-plex Tandem Mass Tag (TMT) reagents (Thermo Fisher Scientific) according to the manufacturer's protocol.

The TMT-labeled samples were mixed equally, desalted, and fractionated on an offline HPLC (Agilent 1220) using basic pH reverse phase liquid chromatography (pH 8.0, XBridge C18 column, 4.6 mm x 25 cm, 3.5 µm particle size, Waters). The fractions were dried, reconstituted in 5% FA plus 0.1% TFA, and analyzed by acidic pH reverse phase C18 LC-MS/MS (New objective, 75 µm ID x ~25 cm, 1.9 µm C18 resin from Dr. Maisch GmbH, heat at 60 $^{o}$C) on an Ultimate 3000 UPLC system (Thermo Fisher Scientific). Peptides were eluted in a 180 min gradient of 15%-40% buffer B (70% ACN, 2.5% DMSO, and 0.1% FA) with buffer A (2.5% DMSO, and 0.1% FA) at a flow rate of ~0.2 µL /min flow rate. The eluted peptides were ionized by electrospray ionization and detected by an in-line Orbitrap Fusion mass spectrometer (Thermo Fisher Scientific) operating in data-dependent (3 s cycle) mode with a survey scan in the Orbitrap (60,000 resolution, scan range 410–1600 m/z, $1 \times 10^6$ AGC target, and 50 ms maximal ion time), followed by sequential isolation of abundant ions in 3 s duty cycle, with fragmentation by higher-energy collisional dissociation (HCD, 38 normalized collision energy), and high resolution detection of MS/MS ions in the Orbitrap (60,000 resolution, $1 \times 10^5$ AGC target, 105 ms maximal ion time,1.0 *m/z* isolation window, and 20 s dynamic exclusion).

MS/MS spectra were searched against UniProt human protein database using the tag-based hybrid search engine, JUMP (*5*). Search parameters included mass tolerance of 20 ppm for precursor ions and MS/MS ions, full trypticity with maximal two missed cleavages, maximal three dynamic modification sites per peptide, dynamic modification with Met oxidation (+15.99492 Da), and static modifications with TMT tags on Lys and N-termini (+229.16293 Da), and Cys carbamidomethylation (+57.02146 Da). All matched MS/MS spectra were filtered by mass accuracy and matching scores to reduce protein FDR to about 1%, based on the target-decoy strategy using a reversed database (*6*).

**Di-GLY TMT ubiquitinome analysis**

Proteomic profiling of ubiquitinome was performed with a previously reported protocol (*8*) with modification. Briefly, for each sample, HEK293T cells from five 15-cm dish were harvested, washed with ice-cold PBS and collected by centrifugation. Cells were lysed in a buffer (50 mM HEPES, pH 8.5, 8 M urea, and 0.5% sodium deoxycholate) with deubiquitinating enzyme (DUB) inhibitors (50 µM PR-619 and 10 mM iodoacetamide) using probe-sonication (15 s × 3 cycles). It should be noted that iodoacetamide-induced pseudo-di-GLY peptides were not detected at room temperature (*9*) and were not recognized by di-GLY antibodies during the enrichment (*10*). To further minimize the impact of DUB activities, the cell lysis protocol was modified by immediately digesting the lysates with LysC (Wako) at an enzyme-to-substrate ratio of 1:100 (w/w) for 2 h at room temperature, to achieve rapid "DUB-deactivation-through-proteolysis."

Following this, the samples were quenched with DTT (30 mM for 30 min) and subjected to trypsin digestion and desalting.

The desalted peptides (about 4 mg) were resuspended in 400 µL ice-cold IAP buffer (50 mM MOPS, pH 7.2, 10 mM sodium phosphate, and 50 mM NaCl), and centrifuged at 21,000 x g for 10 min at 4°C to remove any insoluble material. The peptides were then incubated with di-GLY antibody beads (Cell Signaling Technology) at an antibody-to-peptide ratio of 1:20 (w/w, optimized through a pilot experiment) for 2 h at 4°C with gentle end-over-end rotation. The antibody beads were then collected by a brief centrifugation, washed 3 times with 1 mL ice-cold IAP buffer and twice with 1 mL ice-cold PBS, while the supernatants were carefully removed and saved. The di-GLY peptides were eluted at room temperature twice with 50 µL of 0.15% TFA, dried, resuspended in 50 mM HEPES (pH 8.5) for 11-plex TMT labeling. The TMT labeled peptides were combined equally among replicates, desalted, and analyzed by LC/LC-MS/MS using a similar protocol descried above except the following differences: (i) a 60-min gradient used in the acidic pH LC-MS/MS with Orbitrap QE HF mass spectrometer (Thermo Fisher Scientific), (ii) database search using the COMET algorithm (v2018.013, (*11*)) with 0.02 Da mass tolerance for MS/MS ions, full trypticity with maximal 5 missed cleavages, maximal 5 dynamic modifications per peptide, and di-GLY modification on Lys (+114.04293 Da) as a dynamic modification.

Proteomics statistics

In spectral counting-based quantification experiments (TUBE pulldown and immunoprecipitation), the total number of spectra, namely spectral counts (SCs), matching to individual proteins, reflect their relative abundance in one sample after the protein size is normalized and is useful for comparing the level of the same protein across a number of samples (e.g., control and stressed samples). The summed SC of replicate experiments was compared among different treatments with missing values replaced by a constant low value to calculate fold change and *P* values. The reported *P* value for comparison of SCs for each protein in treated vs. control cells was derived by G-test as described previously (*5*). Stress-induced changes were considered statistically significant with $P \leq 0.02$ and fold change $\geq 2$, excluding proteins detected in only one of three replicates for experiments performed in triplicate. In TMT-based quantification experiments (total proteome and di-GLY profiling), *P* values for comparison between treatments were derived by moderated t-test implemented in limma (R package) (*6*). Stress-induced changes were considered statistically significant *with* $P \leq 0.05$ and fold change $\geq$ 1.5. UpSet plots were generated using UpSetR Shiny App (https://gehlenborglab.shinyapps.io/upsetr/) (*7*) and heat maps were generated using Morpheus (https://software.broadinstitute.org/morpheus). The heat shock ubiquitinome was defined from the TUBE experiment as proteins with statistically significant increase in the soluble fraction and proteins with statistically significant increase in the pellet fraction but not decreased in the soluble fraction (Fig. 2J), excluding 8 proteins for which peptides were measured to have decreased abundance in the di-GLY TMT experiment. To generate the box plot in Figure 3I, the total TMT intensity among all 11 samples for each peptide was summed, the fraction of the total signal for each treatment was calculated, and a box-and-whisker plot for all peptides from heat shock ubiquitinome proteins was generated in Excel. For comparisons in Figures 3J-L, the average abundance as measured by spectral counts of all proteins in the 60-min heat shock TUBE samples were plotted against the total protein abundance as determined by the average TMT intensity in the control samples from the whole proteome analysis (Fig. 3J), the mRNA level as determined by the average read count in control samples from the RNA seq experiment (Fig. 3K), and Riboseq counts from control HEK293T cells determined from a ribosome profiling experiment performed in a previous study (Fig. 3L) (*15*). Data from a previously published study (*16*) was used to generate the histogram of melting temperatures for the heat shock ubiquitinome in comparison to all proteins reported in that study.

RNA-seq

For each sample, HEK293T cells from one 10-cm dish harvested and pelleted by centrifugation. Pellets were washed with ice-cold PBS, and cells were lysed in 600 µl RLT buffer (Qiagen) supplemented with β-mercaptoethanol, and subsequently homogenized by spinning for 1 min at maximum speed in a QiaShredder column (Qiagen). RNA was then extracted according to the manufacturer's protocol using an RNeasy mini kit (Qiagen). RNA was quantified using the Quant-iT RiboGreen assay (Life Technologies) and quality checked by 2100 Bioanalyzer RNA 6000 Nano assay (Agilent) or LabChip RNA Pico Sensitivity assay (PerkinElmer) prior to library generation. Libraries were prepared from 250-1000 ng total RNA using the TruSeq Stranded mRNA Library Prep Kit according to the manufacturer's instructions (Illumina P/N 20020595). Libraries were analyzed for insert size distribution on a 2100 BioAnalyzer High Sensitivity kit (Agilent Technologies) or Caliper LabChip GX DNA High Sensitivity Reagent Kit (PerkinElmer). Libraries were quantified using the Quant-iT PicoGreen dsDNA assay (Life Technologies) or low-pass sequencing with a MiSeq nano kit (Illumina). Sequencing was performed on an Illumina NovaSeq 6000 instrument using paired-end 100 cycle dual indexed chemistry. The reported p value for comparison of stress vs control cells was derived by students t-test.

Sterol analysis

For each sample, HEK293T cells from one 60-mm dish were harvested and pelleted by centrifugation. Pellets were washed twice with ice-cold PBS, frozen, and shipped on dry ice to Creative Proteomics LLC for lipid sterol analysis. Frozen samples were combined with 3 mL ice-cold 75% methanol and 2 ng 19-hydroxy cholesterol (internal standard) per $10^6$ cells. Samples were homogenized on ice using a teflon-coated pestle. 1620 µL chloroform containing 0.01% butylated hydroxytoluene (BHT) was added and samples were vortexed for 120 min at room temperature. Samples were centrifuged for 10 min at 3000 × g and the supernatant was collected to a new tube. The protein pellet was re-extracted, and the extracts were combined with the previous extract from each sample. Extraction solvent was evaporated under nitrogen. Dried lipid extracts were washed 3 times with 10 mM aqueous ammonium bicarbonate, then re-dried under vacuum. Samples were applied to Waters Oasis HLB Prime SPE columns for removal of phospholipids. Column eluant was evaporated and samples were resuspended in methanol containing 0.01% BHT using 20 µl solvent per $10^6$ cells extracted. Sterol species were analyzed by high resolution/accurate mass (HRAM)-LC-MS using a Shimadzu Prominence HPLC coupled to a Thermo Scientific LTQ-Orbitrap Velos mass spectrometer. The LC column was a Phenomenex Synergi HydroRP C18AQ, 2.0 mm × 150 mm, 4 µm, 80 Å LC column with a guard column of matching chemistry. Solvent A was water containing 0.1% formic acid. Solvent B was methanol containing 0.1% formic acid. The flow rate was 250 µL/min and the column oven was held at 50°C. 10 µL of each sample was injected and separated by gradient elution. Column eluent was introduced to the mass spectrometer via a 2-position 6 port valve and an electrospray ionization source. Column eluent was introduced to a Thermo LTQ-Orbitrap Velos mass spectrometer via a heated electrospray ionization source. The mass spectrometer was operated in positive ion mode at 60,000 resolution with full scan MS data collected from 300-700 m/z. Data-depended product ion spectra were collected on the four most abundant ions at 30,000 resolution using the FT analyzer. The electrospray ionization source was maintained at a spray voltage of 4.5 kV with sheath gas at 30 (arbitrary units) and auxillary gas at 10 (arbitrary units). The inlet of the mass spectrometer was held at 350°C, and the S-lense was set to 50%. The heated ESI source was maintained at 300°C. Sterol and oxysterol species were identified as their $[M\text{-}H_2O+H]^+$, $[M\text{-}2H_2O+H]^+$, $[M+H]^+$ and $[M+Na]^+$ ions* under the conditions employed. All sterol species were monitored for four ion types: $[M\text{-}H_2O+H]^+$, $[M\text{-}2H_2O+H]^+$, $[M+H]^+$, and $[M+Na]^+$ and values were combined for total sterol or oxysterol species.

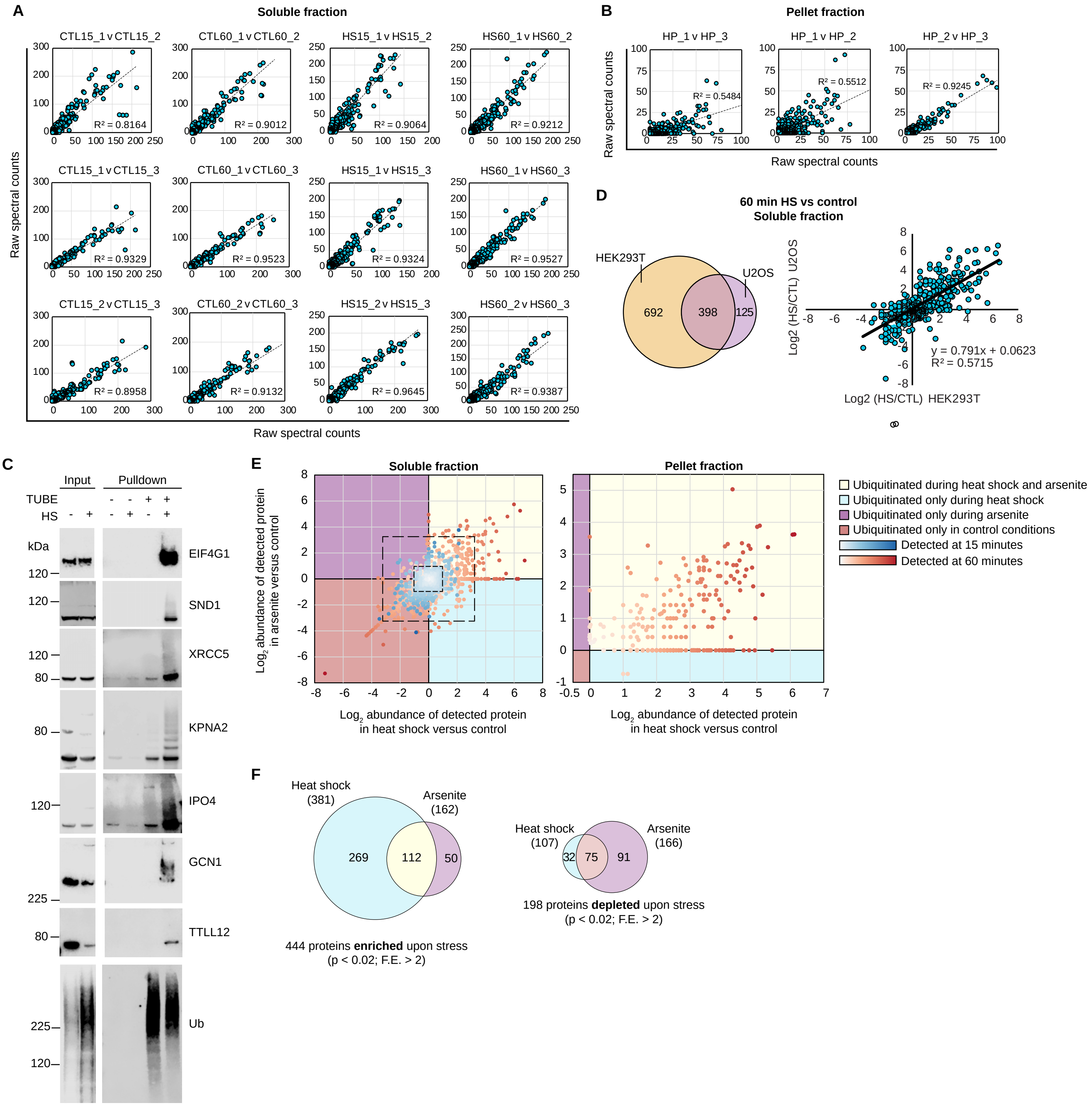


**Fig. S1. Heat shock-induced changes to the ubiquitinome are reproducible, verifiable, not cell type-specific, and share some features with arsenite-induced changes**. (**A**) Correlation of biological replicates of each TUBE soluble fraction sample (see Fig. 2C). (**B**) Correlation of biological replicates of each TUBE pellet fraction sample (see Fig. 2C). (**C**) Immunoblotting was performed to validate results for several proteins detected as having heat shock-dependent increases in ubiquitination in the soluble fraction of the TUBE experiment. In most cases, a smear at a higher molecular weight than the unmodified protein was detected after heat shock, likely indicating modification by poly-ubiquitin. (**D**) Comparison of soluble fraction TUBE experiment results in HEK293T and U2OS cells. The experiment depicted in Figure 2C for 60 min heat shock, soluble fraction samples was repeated using U2OS cells. Overlap of all proteins detected (left) and correlation between heat shock-dependent changes in abundance (right) are shown. (**E**) Correlation between heat shock- and arsenite-dependent changes in the ubiquitinome. Samples for the experiment illustrated in Figure 2C were also collected for cells treated with 0.05 mM sodium arsenite for 15 and 60 min. Correlation between heat shock- and arsenite-dependent changes in abundance (spectral counts) are shown for 15 min (blue dots) and 60 min (red dots) of stress in the soluble (left) and pellet (right) fractions. Color intensity represents fold change, dashed boxes represent 2- and 10-fold change in abundance. (**F**) Overlap of statistically significant increases (top) and decreases (bottom) in ubiquitination determined from the TUBE experiment.

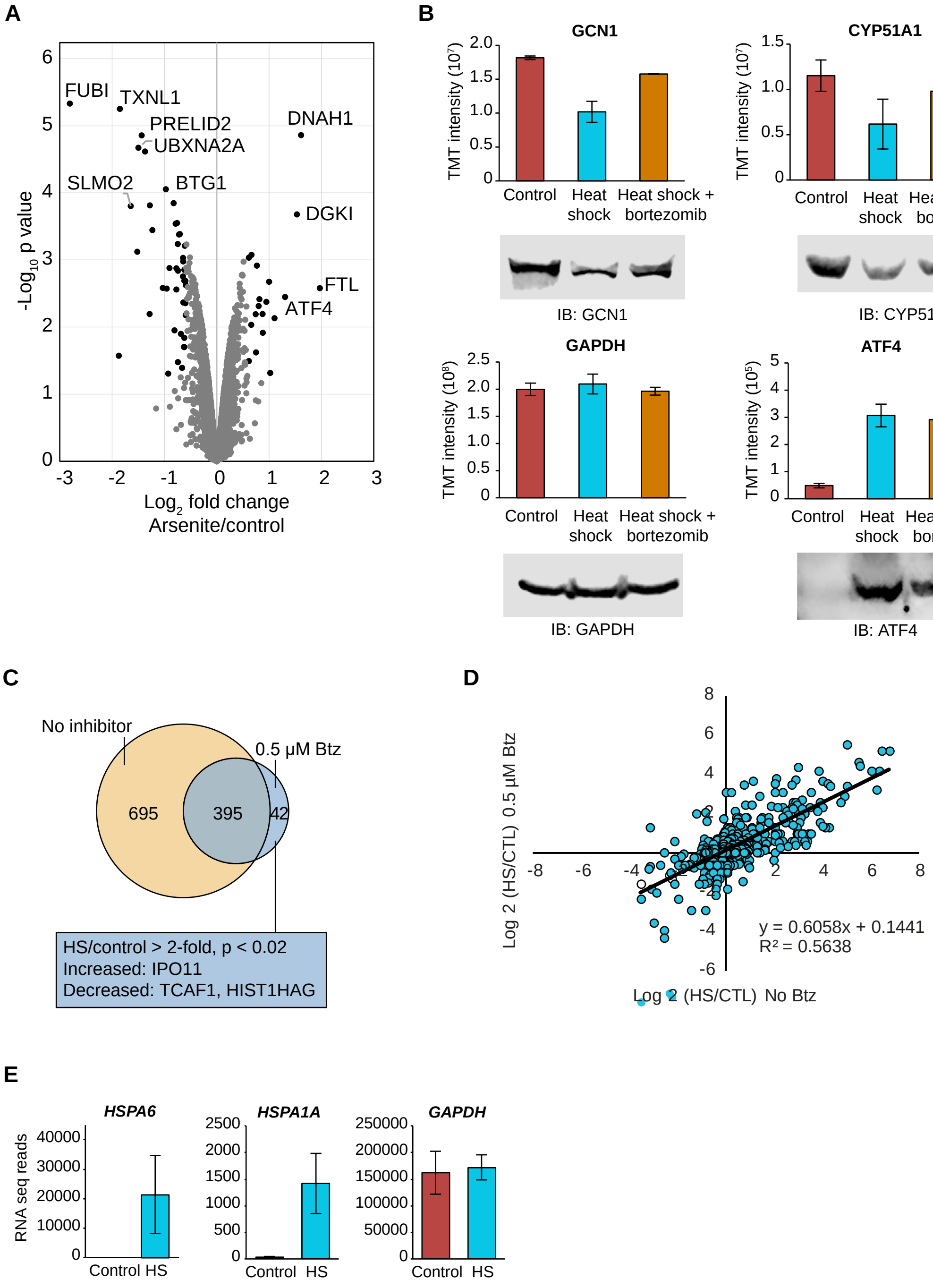


**Fig. S2. Total proteomic and transcriptomic analysis and treatment with proteasome inhibitors reveal additional details of heat shock and arsenite ubiquitinomes.** (**A**) Volcano plot indicating changes in total protein abundance for arsenite versus control samples. (**B**) TMT intensity abundance values and immunoblot validation are shown for representative proteins. Error bars indicate s.d. (**C**) Overlap of statistically significantly heat shock-dependent increased proteins in TUBE experiments in presence or absence of bortezomib (Btz). Blue box lists all proteins with statistically significant changes that were detected only in cells treated with bortezomib. (**D**) Correlation of heat shock-dependent changes in abundance in TUBE experiments in the presence or absence of 0.5 µM bortezomib. (**E**) Raw RNA-seq read count for two notable stress responsive chaperone genes and GAPDH are shown. Error bars indicate s.d.

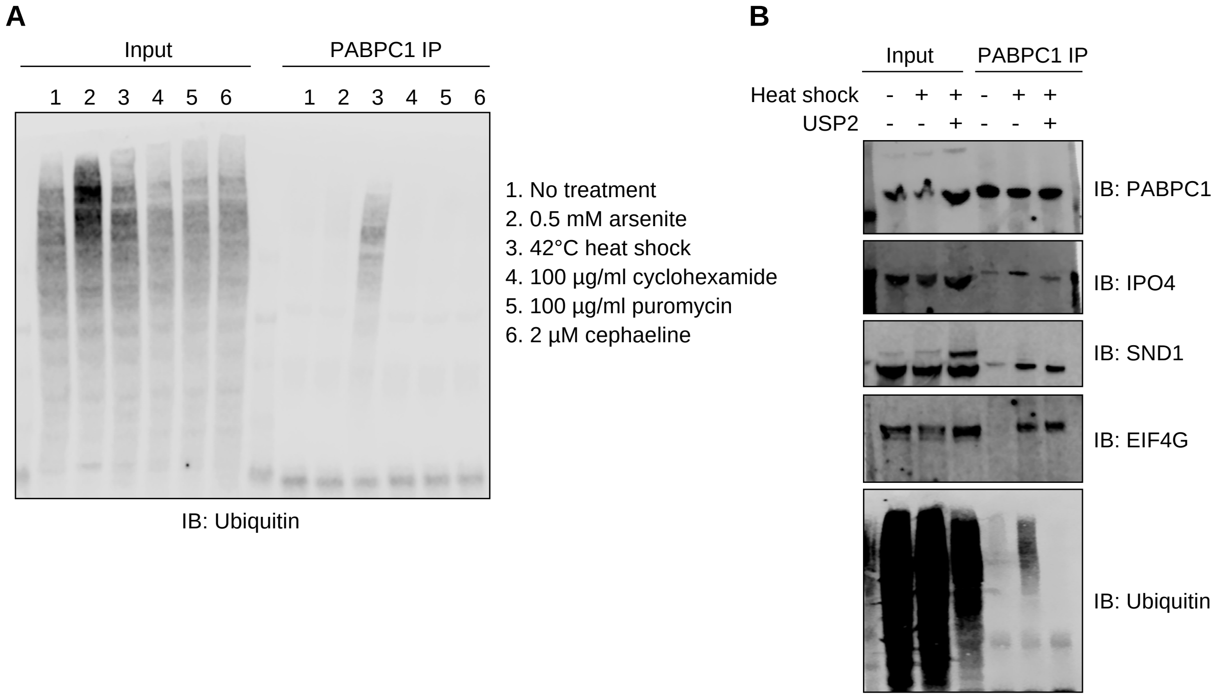


**Fig. S3. Stress-induced ubiquitination of mRNP complexes is specific to heat stress; complex formation is resistant to RNase and deubiquitinase treatment, but requires active ubiquitination for recovery.** (**A**) PABPC1 immunoprecipitation was performed on lysates from cells treated as indicated for 60 min prior to lysis. Input and IP fractions were immunoblotted for poly-ubiquitin. (**B**) PABPC1 immunoprecipitation was performed on lysates before and after heat shock in the presence or absence of deubiquitinating enzyme Usp2 added prior to incubation with antibody-conjugated beads.

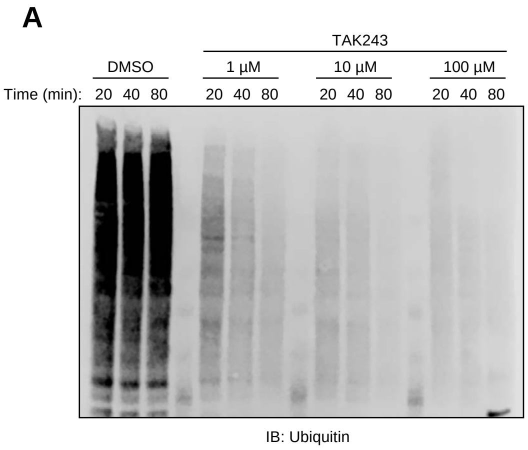


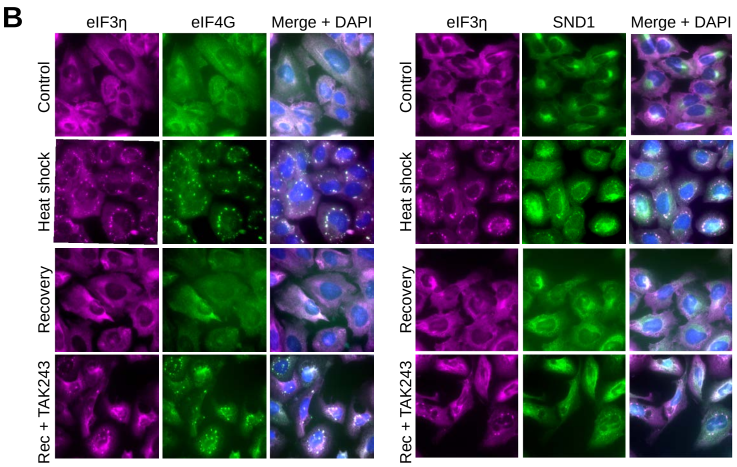


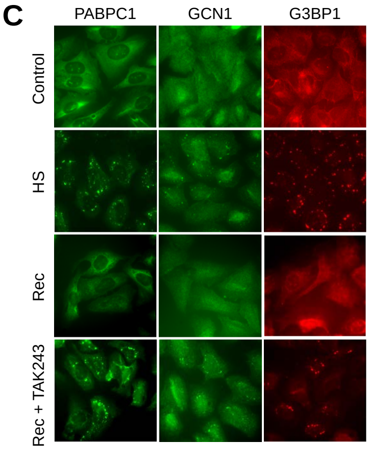


D

**HS and TAK243 puncta formation of endogenous proteins (IF)**

| Protein name | Baseline | Heat shock | Recovery | Recovery + TAK243 |
|---|---|---|---|---|
| PABPC1 | - | + | - | + |
| EIF3eta | - | + | - | + |
| EIF4G1 | - | + | - | + |
| G3BP1 | - | + | - | + |
| SND1 | - | + | - | + |
| GCN1 | - | + | - | + |
| KPNA2 | - | + | - | + |
| SFPQ | - | + | - | + |
| XRCC5 | - | + | - | + |
| HNRNPA2B1 | - | + | - | + |
| CYP51A1 | - | - | - | - |
| EPRS | - | - | - | - |

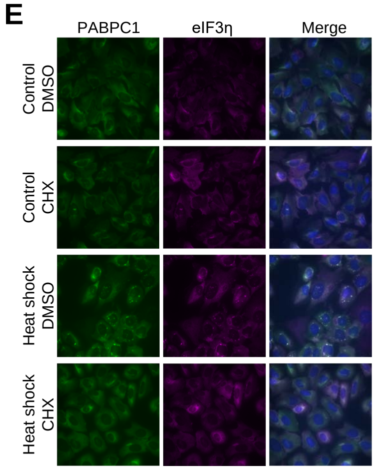


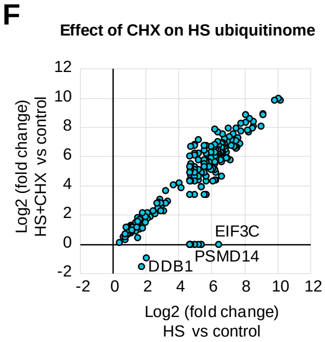


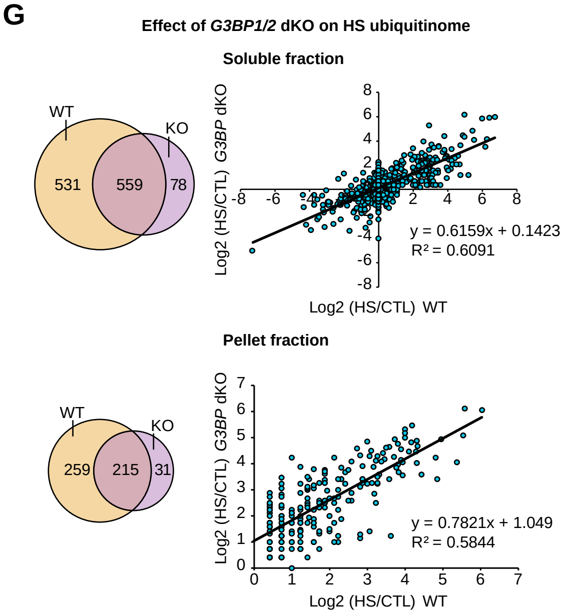

**Fig. S4. Immunostaining of endogenous proteins confirms that heat shock-induced stress granules require active ubiquitination for disassembly, whereas stress granule formation is not a prerequisite for heat shock-dependent ubiquitination.** (**A**) Immunoblot verification of time- and dose-dependent depletion of cellular poly-ubiquitin conjugates in cell lysates following addition of TAK243 directly to the culture media as indicated prior to lysis. (**B**) U2OS cells were fixed and stained for stress granule marker EIF3η and indicated heat shock ubiquitinome proteins after no stress, 60 min heat shock, or 60 min heat shock with 90 min recovery in the presence or absence of 1 µM TAK243. (**C**) Additional results from immunofluorescence imaging of endogenous proteins treated as described in (A). (**D**) Summary of immunofluorescence imaging results including those in (B) and (C) and those for several additional proteins. (**E**) U2OS cells were fixed and stained for stress granule markers PABPC1 and EIF3η after no stress or 60 min heat shock in the presence or absence of 100 µg/ml cycloheximide. (**F**) Correlation of the heat shock ubiquitinome in presence or absence of 100 µg/ml cycloheximide. Untreated and cycloheximide-treated cells were collected and analyzed in parallel in a separate experiment from the dataset illustrated in Figure 2C and correlation of heat shock-dependent changes for statistically significantly increased proteins is shown. (**G**) Comparison of TUBE experiment results in wild-type and *G3BP1/2* double KO HEK293T cells. The experiment depicted in Figure 2C (for 60 min heat shock) was repeated using double KO cells and compared to the previously generated dataset shown in Figure 2C. Overlap of all proteins detected (left) and correlation between heat shock-dependent changes in abundance are shown.

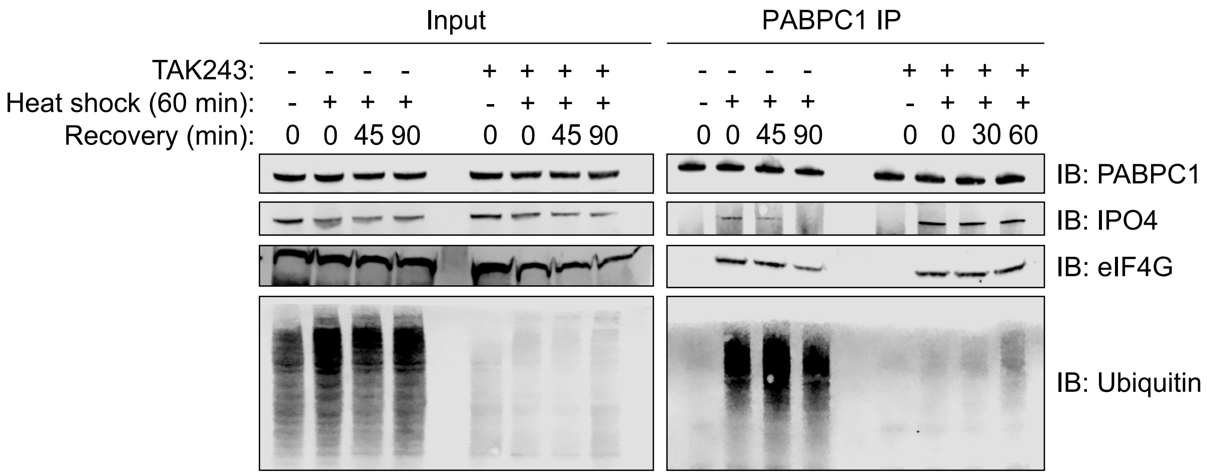


**Fig. S5. Immunoblot showing the requirement of ubiquitination for the reversal of mRNP remodeling.** Immunoblot analysis of formation and dissolution of poly-ubiquitinated protein-mRNA complexes formed during heat shock and isolated by PABPC1 immunoprecipitation.

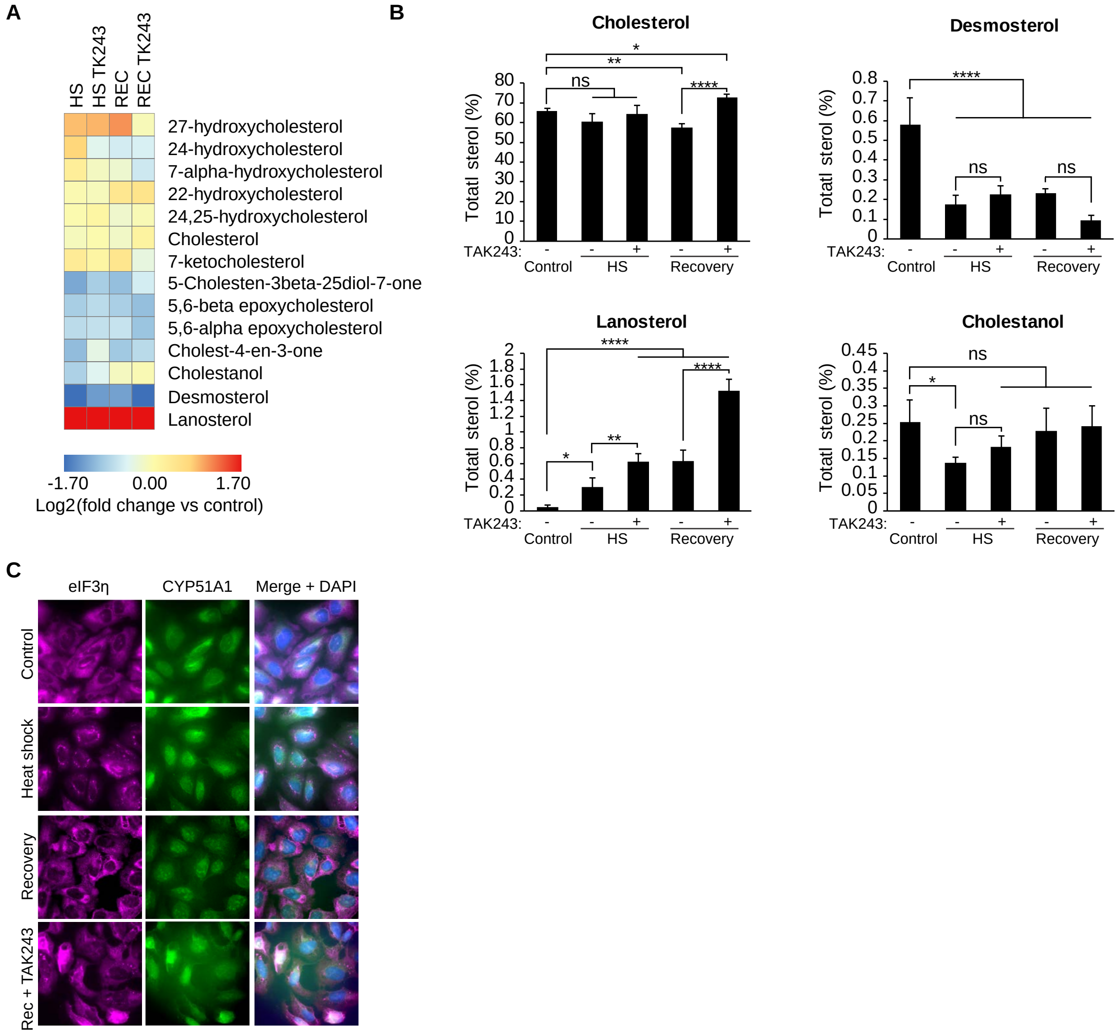


**Fig. S6. Inhibition of ubiquitination effects the cholesterol biosynthesis pathway both during heat shock and recovery in a pattern different than observed for stress granule dynamics, mRNP remodeling, nucleocytoplasmic transport and translation.** (**A**) Heatmap illustrating changes in abundance in sterol metabolites during heat shock and recovery in the presence or absence of TAK243. (**B**) Abundance is shown for several representative sterols in each experimental condition. Graphs represent the average and s.d. of four biological replicates. ns, not significant; **P* < 0.05, ***P* < 0.01, ****P* < 0.001, *****P* < 0.0001, ANOVA with Tukey's test. (**C**) U2OS cells were fixed and stained for stress granule marker EIF3η and cholesterol metabolism enzyme CYP51A1 after no stress, 60 min heat shock, or 60 min heat shock with 90 min recovery in the presence or absence of 1 µM TAK243, indicating CYP51 does not colocalize with stress granules.

**Movie S1.**
Live cell imaging of DMSO control-treated U2OS cells stably expressing GFP-G3BP1 during arsenite stress and recovery.

**Movie S2.**
Live cell imaging of TAK243-treated U2OS cells stably expressing GFP-G3BP1 during arsenite stress and recovery.

**Movie S3.**
Live cell imaging of DMSO control-treated U2OS cells stably expressing GFP-G3BP1 during heat shock and recovery.

**Movie S4.**
Live cell imaging of TAK243-treated U2OS cells stably expressing GFP-G3BP1 during heat shock and recovery.

**Table S1. (separate file) Stress ubiquitinome pilot data.** Table showing results from TUBE-proteomic analysis of cells treated with the five stresses listed in Figure 1B.

**Table S2. (separate file) Heat shock-induced changes in ubiquitination.** Table showing spectral counting results from TUBE analysis of cells treated as described in Figure 2C, including calculated *P* values and fold change for statistically significant changes.

**Table S3. (separate file) Additional TUBE results.** Table showing spectral counting results of TUBE experiments from additional cell lines and cells treated with chemical inhibitors.

**Table S4. (separate file) Heat shock-induced changes in ubiquitination.** Table showing spectral counting results from TUBE analysis of cells treated with arsenite stress, including calculated *P* values and fold change for statistically significant changes.

**Table S5. (separate file) Stress dependent-changes to the total proteome.** Table showing TMT intensity for total proteome analysis.

**Table S6. (separate file) Heat shock-dependent changes to transcriptome.** Table showing read counts from RNA-seq transcriptome analysis.

**Table S7. (separate file) Heat shock-induced changes in di-GLY profiling.** Table showing TMT intensity for di-GLY profiling.

**Table S8. (separate file) Heat shock and arsenite ubiquitinomes.** List of proteins with heat shock- and arsenite-induced statistically significant increases in ubiquitination determined by

TUBE-based proteomics, including correction based on di-GLY analysis for the heat shock ubiquitinome.

**Table S9. (separate file) Heat shock-induced changes in mRNP complex interactors.** Table showing spectral counting results from immunoprecipitation-based proteomic analysis described in Figure 5.

**Table S10. (separate file) Stress granule proteome.** Table showing list of proteins in the stress granule proteome and the high-confidence stress granule proteome.

### References and Notes


1. M. D. Figley, G. Bieri, R. M. Kolaitis, J. P. Taylor, A. D. Gitler, Profilin 1 associates with stress granules and ALS-linked mutations alter stress granule dynamics. *J Neurosci* **34**, 8083-8097 (2014).
2. S. Berg *et al.*, ilastik: interactive machine learning for (bio)image analysis. *Nat Methods* **16**, 1226-1232 (2019).
3. C. McQuin *et al.*, CellProfiler 3.0: Next-generation image processing for biology. *PLoS Biol* **16**, e2005970 (2018).
4. P. Xu, D. M. Duong, J. Peng, Systematical optimization of reverse-phase chromatography for shotgun proteomics. *J Proteome Res* **8**, 3944-3950 (2009).
5. J. Y. Zhou *et al.*, Galectin-3 is a candidate biomarker for amyotrophic lateral sclerosis: discovery by a proteomics approach. *J Proteome Res* **9**, 5133-5141 (2010).
6. G. K. Smyth, Linear models and empirical bayes methods for assessing differential expression in microarray experiments. *Stat Appl Genet Mol Biol* **3**, Article3 (2004).
7. A. Lex, N. Gehlenborg, H. Strobelt, R. Vuillemot, H. Pfister, UpSet: Visualization of Intersecting Sets. *IEEE Trans Vis Comput Graph* **20**, 1983-1992 (2014).
8. Y. A. Valentin-Vega *et al.*, Cancer-associated DDX3X mutations drive stress granule assembly and impair global translation. *Sci Rep* **6**, 25996 (2016).
9. P. Leuenberger *et al.*, Cell-wide analysis of protein thermal unfolding reveals determinants of thermostability. *Science* **355**,  (2017).
